# Hyper-dominant species drive unexpected functional shifts in North American birds

**DOI:** 10.1101/2025.07.23.666479

**Authors:** Heng-Xing Zou, Adam C. Smith, Yiluan Song, Kai Zhu, Brian C. Weeks

**Affiliations:** Institute for Global Change Biology, University of Michigan, Ann Arbor, MI 48109; School for Environment and Sustainability, University of Michigan, Ann Arbor, MI 48109; Department of Biology, Carleton University, Ottawa K1S 5B6, ON, Canada; Canadian Wildlife Service, National Wildlife Research Centre, Environment and Climate Change Canada, Ottawa K1S 5R2, ON, Canada; Michigan Institute for Data and AI in Society, University of Michigan, Ann Arbor, MI 48109

## Abstract

Global environmental changes have led to widespread intraspecific shifts in functional traits across large numbers of widely distributed taxa. However, how within-species functional shifts predict community-level changes is unknown. We summarized seven key traits that are hypothesized to shift adaptively under environmental changes using local community-weighted means across North American bird communities. We find that communities shifted toward larger body mass and smaller appendage sizes over time, the opposite of widely observed within-species temporal changes in North American birds. These unexpected patterns arise from changes in the relative abundance of a very small number of hyper-dominant species, with 23 out of 470 (5%) species explaining the community-level trajectories in functional traits. These species are known to be heavily impacted—both positively and negatively—by human activities, and the idiosyncratic drivers of population sizes result in weak associations between community-level functional shifts and major bioclimatic variables but strong associations with human-induced land use changes. Our results illustrate the complexity in scaling from within-species responses to global change to community-level responses and highlight the importance of species-specific responses to the environment beyond climate when predicting functional and taxonomic diversity under global change.

## Main Text

Rapid climate change has driven widespread shifts in species’ abundances and distributions (1–3), as well as changes in key functional or life history traits (4, 5). Both shifts in species abundances across space and functional traits through time are expected to alter the taxonomic and functional diversity of ecological communities (6, 7). Global change drivers are often general, such as broadly consistent increases in temperature and widespread human-mediated habitat degradation (8, 9). Many of these changes could lead to similarly consistent responses among taxa; for example, as the world warms, thermoregulatory demands are likely to shift similarly across broad groups of species, driving parallel adaptations to facilitate heat dissipation (10). However, impacts of other global change factors, such as land use change, could be more idiosyncratic among species, potentially resulting in inconsistent responses (e.g., variable shifts in range (11), phenology (12), and morphology (13) across species). These inconsistencies may pose remarkable challenges for predicting how ecological communities will respond to global change from within-species responses.

Birds are a model system for relating species- and community-level responses to global change as they have comprehensive distributional (14), phylogenetic (15), behavioral (16), and functional trait (17) data, as well as long-term time series data on shifts in range, phenology, and morphology. There is reason to expect that responses to rapid environmental changes are general and directional among avian communities. Within species, shifts in key functional traits have been relatively uniform: broadly, species have become smaller (4, 18–20) and their appendages, such as beaks, tarsi, and wings, have become longer (13). These shifts are often interpreted within the context of thermoregulation or adaptation to more fragmented habitats and urbanization (10, 21, 22). There is growing evidence that some within-species responses to climate change (e.g., range shifts) have resulted in community-level changes. Bird species have generally shifted poleward and up in elevation, likely in response to the warming climate (2, 12, 23), and these distributional changes are thought to have contributed to consistent local changes in functional richness and evenness across large scales (7, 24, 25).

Should within-species changes in morphology represent consistent selection favoring particular phenotypes (e.g., smaller body size) under changing climate and environments, we may expect that species with that phenotype should be increasing in relative abundance, shifting the community-weighted mean (CWM) of that trait in parallel with the intra-specific trends. Alternatively, scaling up within-species trends to the community level may be more complex. Idiosyncratic responses of individual species to the broad array of environmental changes that are occurring contemporaneously may further lead to inconsistencies between shared within-species responses to a single driver (e.g., climate change) and community-level functional shifts (20). Relatedly, within-species changes may be adaptive, but this micro-evolutionary response may not predict variation in fitness among species along the same trait axis. Finally, while many within-species changes have been associated with environmental change, it is unclear if these changes are adaptive or the result of plasticity, in which case they may be less likely to predict shifts in fitness among species. Evaluating the generality of these functional responses across the scales is important in predicting shifts in bird community structure and ecosystem function under rapid environmental changes.

### Unexpected community-level shifts in functional traits

To test if community-level functional shifts align with previous hypotheses based on within-species changes, we characterized changes in functional traits of local communities based on abundance data for 470 species spanning half a century of sampling in North America (North American Breeding Bird Survey, NABBS; Fig. 1A; (26)). To obtain robust estimates of local community composition, we aggregated abundance data within one-degree grids, aligning with the design of NABBS to have at least one route per grid with suitable access roads (see Methods). We calculated community-weighted means (CWMs) of seven key morphological and demographic traits: beak size, wing size, body mass, relative beak length, relative wing length, clutch size, and generation length. We calculated corrected beak and wing lengths (Methods); like relative beak/wing lengths, these measurements correct for the associations between beak/wing length and body mass (27). Similarly, we also corrected clutch size and generation lengths by body mass (Methods). CWMs are the average of functional trait values among species within a community, weighted by species abundances (28) and provide more mechanistic interpretations of functional shifts at the community level, more direct comparisons with within-species shifts, and are less biased by the presence of rare species than ordination-based approaches (29). To better compare with within-species expectations of functional shifts, we examined how shifts in CWMs are associated with time and environmental variables. We considered major climatic attributes and human-induced land use changes, which have greatly affected ecological communities (9, 30, 31).

**Fig. 1.**
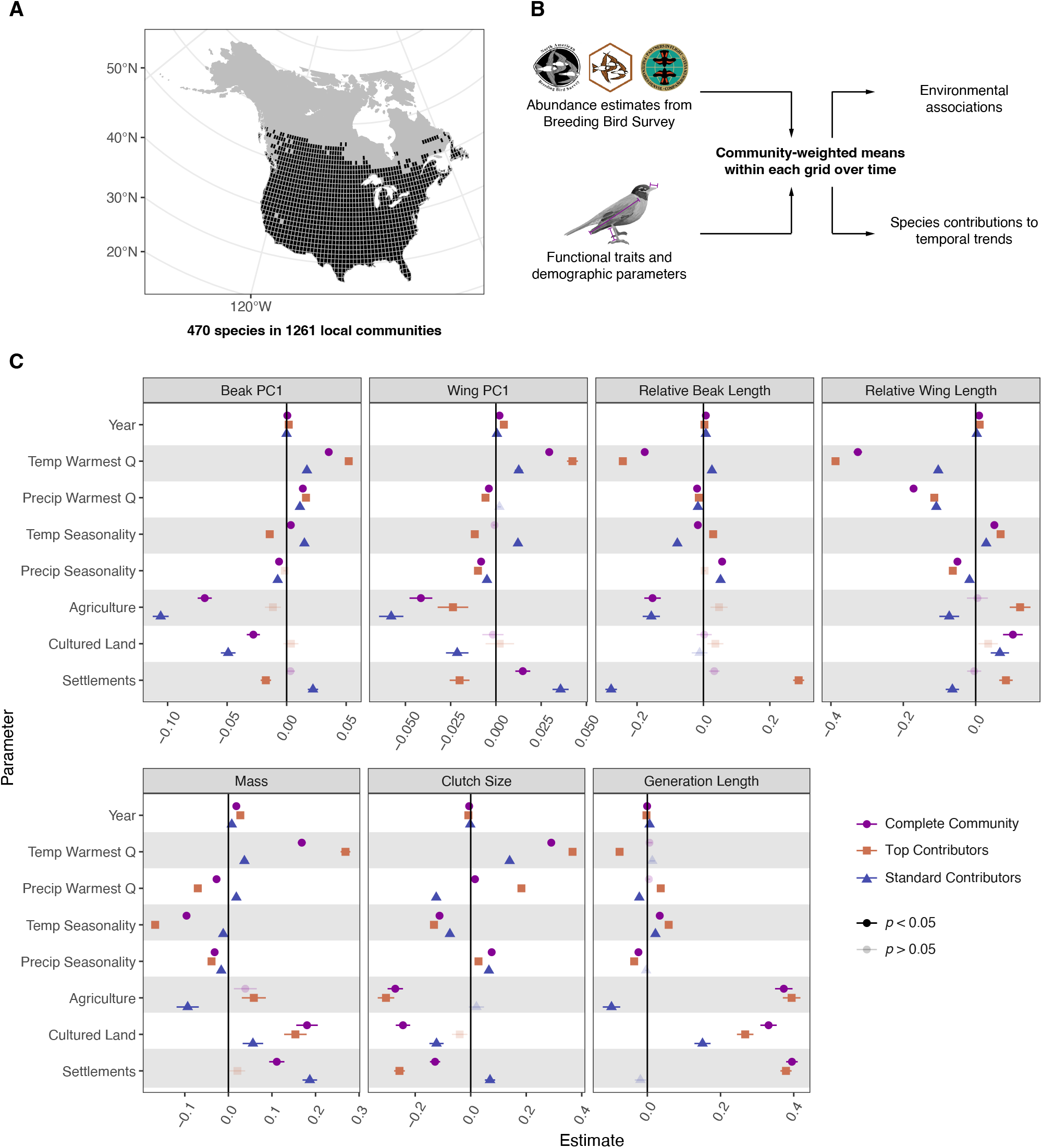
Overview of the analyses and associations between functional shifts and environmental variables. **A**. Map of the study region, showing the 1261 local communities (one by one degree grids) included in the analyses. **B**. Overview of the analyses. **C**. Associations between community-weighted means (CWMs) of seven key functional traits and main environmental variables, including climate, human-induced land use, and year. CWMs are calculated from either all species in the communities (complete communities), only the top contributors, or only the rest of the species (standard contributors). All statistics are shown in table S1. Illustration by Olivia Stein.

We find that community-level shifts in key morphological traits are often in opposite directions from within-species expectations based on adaptations to changing climates. CWM of body mass increases with temperature of the warmest quarter (mean = 0.168, standard error = 0.00893, *p* < 0.0001; Fig. 1C), contrary to within-species studies that have found reductions in body size as the climate has warmed over time (4, 19). The CWMs of beak and wing sizes (the first PC axes from phylogenetic PCA of four beak measurements and three wing measurements, respectively; see Methods) also positively associate with temperature (beak: 0.0354 ± 0.00177; wing: 0.0295 ± 0.00205; both *p* < 0.0001), but relative beak and wing lengths both show strongly negative associations with temperature (beak: -0.177 ± 0.00868; wing: -0.326 ± 0.0117; both *p* < 0.0001). These results contrast with within-species trends toward longer relative appendage size (4, 19, 20) that are often attributed to selection towards increased effective surface area and heat dissipation efficiency under warming (10). The CWM of clutch size positively associates with temperature (0.290 ± 0.00980, *p* < 0.0001) but negatively associates with temperature seasonality (-0.112 ± 0.00480, *p* < 0.0001; Fig. 1C). Both are contrary to within-species patterns: higher ambient temperature is associated with smaller clutch sizes often attributed to increased potential for hatching failure, and higher intra-annual fluctuations in temperature may select for larger clutch sizes as a bet-hedging strategy (32–34). Generation length represents the average age of parents in any population of a species (35, 36) and higher intra-annual fluctuations in temperature should select for earlier reproduction (35, 37). In contrast, we observed a positive association with temperature seasonality (0.0339 ± 0.00393, *p* < 0.0001). Because temperature and precipitation seasonality only directly affect year-round residents (35), we re-analyzed the associations between climate seasonality and demographic traits separately for species that undertake long-distance seasonal migrations and those that do not according to (38). The signs of all associations are consistent among migrants and residents, but the positive association between the CWM of generation length and temperature seasonality is non-significant (fig. S1-S2).

Many studies have also suggested human-induced land use changes should favor species with certain traits, especially in urban areas where habitats are most modified (21, 22). We found that these land use changes often have stronger associations with community-level functional traits, but some of them are again contrary to expectations. The CWM of body mass is positively associated with human-modified forests and drylands (cultured land), and villages and urban areas (settlements; Fig. 1C; table S1). These results are different from the expectation that smaller body size should be favored in more human-dominated landscapes such as urban areas due to lower resource and habitat requirements (21, 22). The CWM of relative wing length is positively associated with cultured land (0.103 ± 0.00272, *p* = 0.000196) but not agricultural land and dense settlements, again different from the expectation that urban-adapted species should have larger wing-to-body ratios to assist with dispersal among fragmented habitats (21, 22). All three human-induced land use changes are negatively associated with the CWM of clutch size (agriculture: -0.272 ± 0.0277, cultured: -0.244 ± 0.0259, settlements: -0.128 ± 0.0184; all *p* < 0.0001), contrary to the expectations that anthropogenic environmental stress should favor species with larger clutch sizes (21, 22). On the other hand, the CWM of generation length is positively associated with all three human-induced land use changes (agriculture: 0.373 ± 0.0239, cultured: 0.331 ± 0.0223, settlements: 0.395 ± 0.0157; all *p* < 0.0001). Because maximum longevity has a strongly positive correlation with generation length (35, 36), our result is therefore consistent to the expectation from previous studies that longer-lived species are favored in urban areas (21, 22).

We found that across North America, community-level shifts in morphological and demographic traits often contrast with species- and population-level expectations. We did not consider how species traits may change across space or time, which is unlikely to alter results because the magnitudes of within-species trait changes are much smaller than typical among-species variation in these traits (39). In addition to advancing our understanding of how communities have shifted through time, our findings shed new light on the interpretation of well-documented functional shifts within species. If these changes in morphological and demographic traits are adaptive to environmental changes, then it is intuitive to expect that species with traits in these more adaptive directions would increase in abundance and lead to shifts in the CWMs of the trait that are aligned with intra-specific trends. However, the shifts we find at the community level are often contrary to the shifts that have been broadly observed within species, which we interpret in three possible ways. First, observed within-species functional shifts may not be adaptive, or may not have meaningful effects on fitness (39–41). Indeed, although some morphological responses are consistent over time, whether they are indeed adaptive to environmental changes remains unclear (39, 40). With more studies finding evidence both supporting and opposing longstanding hypotheses based on the adaptation of climate and environmental change (e.g., warming and urbanization (10, 22)), the lack of consistency on a community level in our results may further indicate that within-species morphological changes may not be responding to the same single driver in the environment. Second, if observed intraspecific functional shifts are indeed adaptive, then our results indicate that microevolutionary processes may not necessarily predict relative fitness differences among species (42). Third, factors that drive adaptive intraspecific changes in functional traits (e.g., climate change) may have weaker effects on relative abundance than other drivers that do not drive intraspecific trait evolution but do determine differential sensitivity to more significant population-level pressures (e.g., human-induced land use change; (43)). As such, inconsistencies between intraspecific trait changes and shifts at the community level demonstrate the complexities involved in scaling up from species to community level dynamics in the context of multifaceted global environmental change.

### Nearly universal functional shifts across space

Continental-scale analyses of functional shifts may mask local and regional environmental factors. We explored the generality of functional shifts across local communities in North America by analyzing the temporal trajectory of each local community within a functional space constructed by a PCA that reduced the CWMs of the seven traits into two major axes (Fig. 2A). The first PC axis (which explains 43.9% of the variance) mostly represents variation in traits related to absolute body size (beak and wing sizes and body mass); the second PC axis (which explains 16.9% of the variance) mostly reflects variation in clutch size, generation length, and relative wing length (Fig. 2A). We focus on the locations of the community in functional trait space at the start and end years (1970 and 2021), summarized by their positions in quadrants, and the overall direction (angle) of the shift for each community, summarized by the quadrant it points to (Fig. 2A). In 1970, most local communities in central North America were characterized by larger body mass, larger absolute appendage sizes, larger clutch size, and shorter generation lengths (Fig. 2B; fig. S3). In 2021, many local communities, especially at lower latitudes, were characterized by smaller clutch sizes and longer generation lengths. Examining the directions of these shifts from 1970 to 2021, we observe an east-west divide, with local communities in the Eastern Forests and eastern parts of the Prairie regions shifting towards smaller clutch sizes and longer relative beak and wing lengths, whereas many other major avifaunal biomes in the west and north have shifted towards larger body mass and clutch sizes (Fig. 2B; fig. S4; see fig. S5 for a map of avifaunal biomes). Overall, these local-level functional shifts are broadly consistent with the results from the continent-level trends, that CWM of body mass have generally increased with time and temperature, and that clutch size and generation lengths have generally decreased with human-induced land use changes (Fig. 1C). The highly consistent community-level functional shifts that contrast with within-species expectations highlight that even when there are nearly universal changes in the environment over time that can drive consistent intraspecific functional trends among species (e.g. consistent size declines associated with increasing temperatures; (4)), these conserved trends may not scale simply to analogous functional shifts of local communities.

**Fig. 2.**
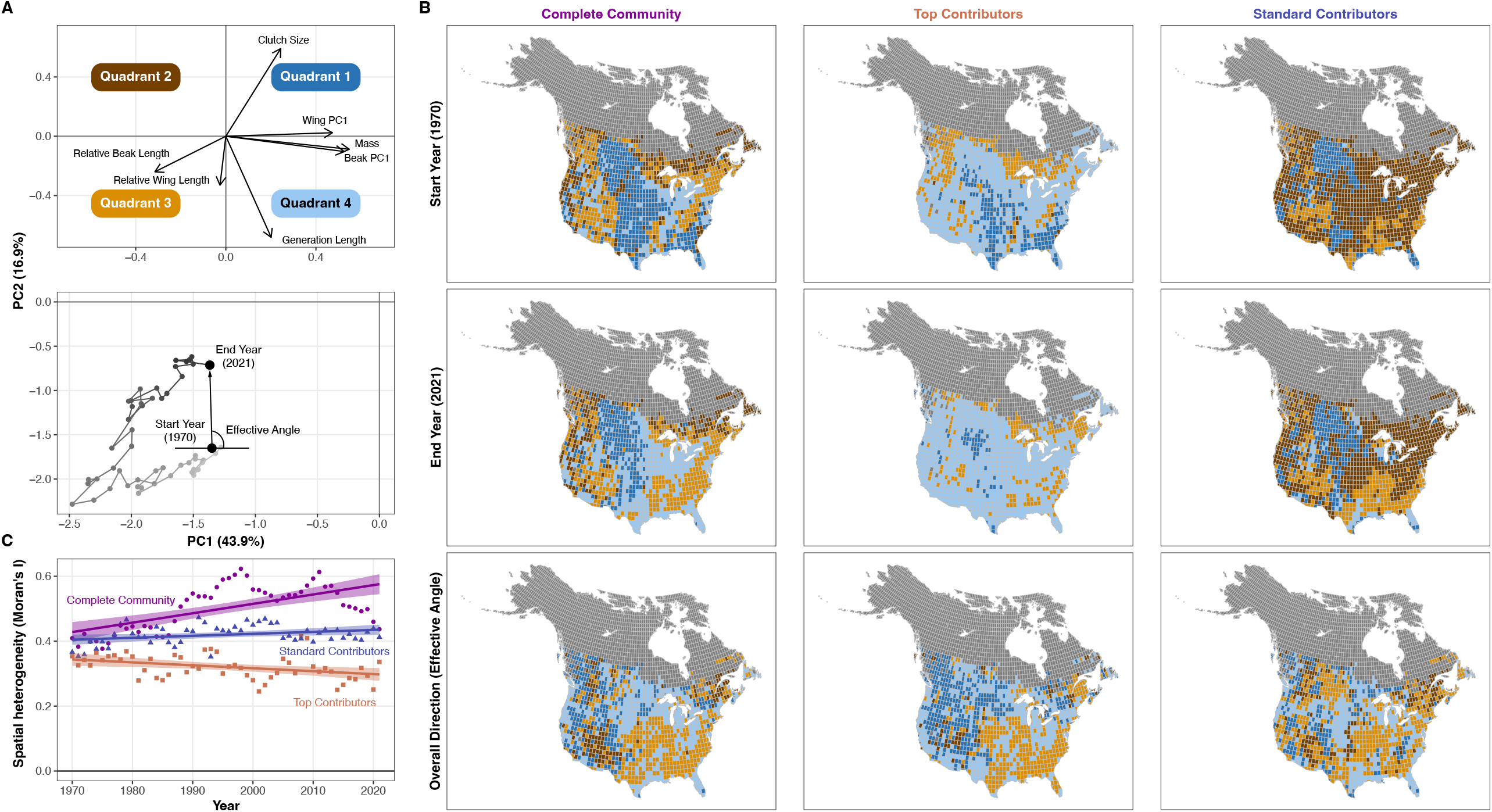
Analyzing the temporal trajectory of local communities in the functional space. **A**. The functional space constructed by PCA of all communities by their community-weighted mean measurements over time, and a temporal trajectory of one local community (40°N, 110°W) moving through the functional space over time, showing the start and end points and the overall direction (effective angle). In the trajectory, lighter shades represent earlier years, and darker shades represent later years. The two panels represent the same functional space, but the scales are adjusted for better visualization. **B**. Maps of each local community’s start and end positions in the functional space, measured by the quadrants in **A**, and the overall direction of the functional shifts, measured by the quadrants to which the trajectory travels. Columns show results for complete communities, subcommunities with top contributors, and subcommunities with the standard contributors. **C**. Spatial heterogeneity of functional positions of the base communities, top contributors, and the rest of the species, calculated as Moran’s *I* from the maps. A Moran’s *I* closer to 0 means lower spatial heterogeneity.

### Functional shifts driven by a few hyper-dominant species

As a consequence of global environmental changes, shifts in the composition of a community may emerge due to consistent changes across many of the species in a community (e.g. consistent declines in the abundances of smaller species, resulting in an increase in the CWM of mass through time) or extreme shifts in the abundance of a relatively small number of species that either become hyper-dominant while others are extirpated or decline rapidly transitioning away from being a hyper-dominant species (44). To better understand the mechanisms underlying the unexpected functional shifts on the community level, we quantified the contributions of each species to the temporal trends of CWMs within each local community, allowing us to determine the magnitude of the contributions of individual species to community-level functional shifts. For each trait, we ranked the species by the number of local communities in which they contribute greatly to the temporal trend (i.e., they are within the top 10% of species driving the community trend in that local community). We found 4.89% of species (23 out of 470) had overwhelmingly important contributions to functional shifts across North America (Fig. 3; fig. S6-S12). To illustrate the importance of these top contributors to community-level functional shifts, we calculated CWMs for two subcommunities: (1) only the “top contributors”, and (2) the rest of the species (hereafter the “standard contributors”). We then repeated all the analyses of the temporal trends and environmental associations with CWMs and compared the results of each subcommunity with the results derived using the full set of species (the “complete communities”).

**Fig. 3.**
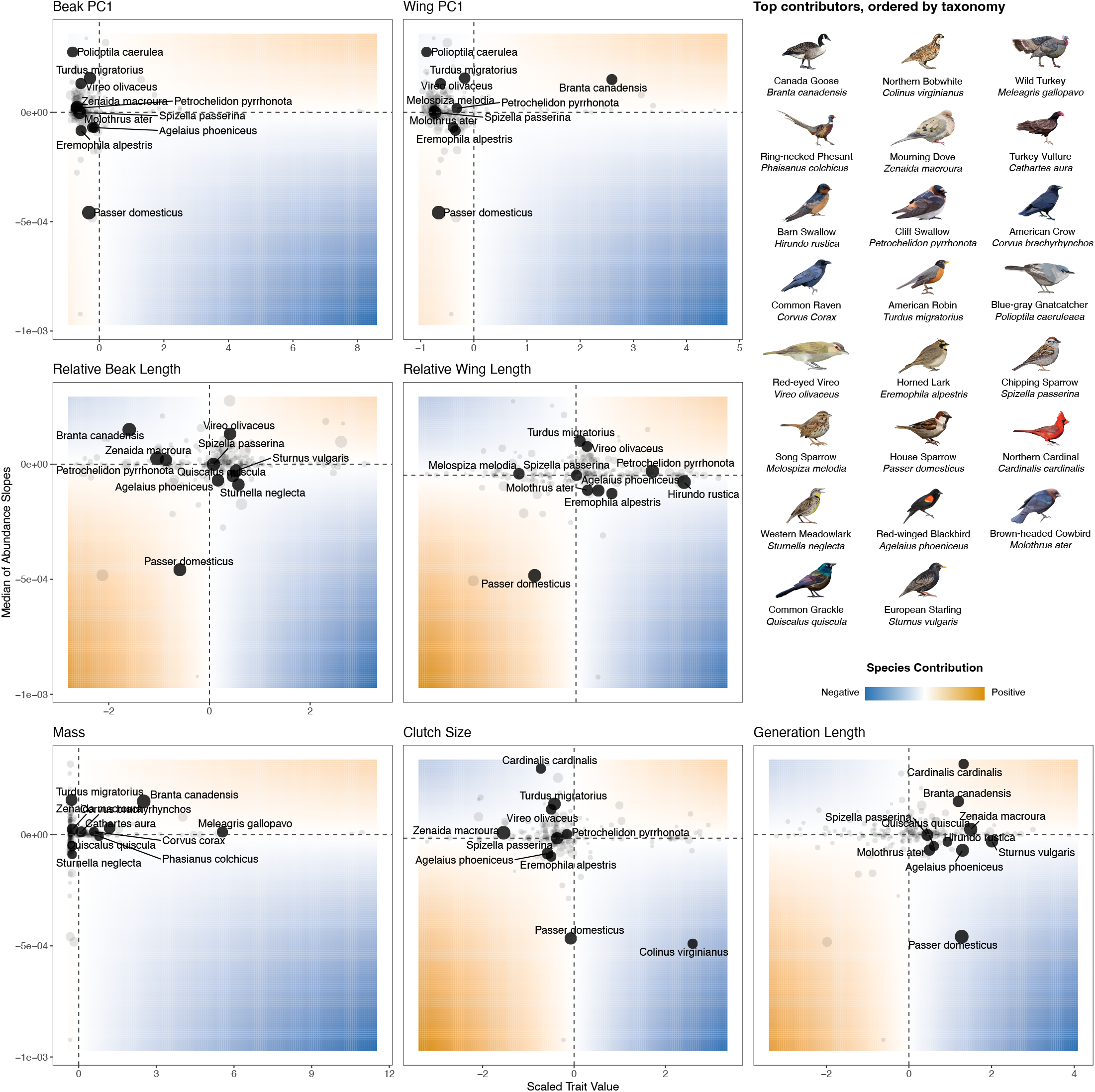
Species contributions to the community-weighted means (CWMs) of seven key functional traits. Panels only include species that contribute substantially (ranked top 10% of all species in a local community) in at least one local community; point size denotes the number of grids in which a species contributes greatly. The sign of the contribution depends on the species’ temporal trends of relative abundance and its trait value, e.g., if a species with below-average trait value increases in relative abundance, it will contribute negatively by lowering the CWM of the trait. For each trait, the identity of the top 10 species that contribute to the CWMs of the most local communities is labeled; pooling all these species results in 25 top contributors of the observed functional shifts over time, shown on the right. For each panel, the x-axis is the scaled trait values, and the y-axis is the median of the temporal slopes of relative abundances across each species’ range. Shades denote the absolute magnitudes of species contributions: colder colors represent negative contributions, and warmer colors represent positive contributions. Axes and scales are different for each panel for better visualization. Illustrations by Olivia Stein.

In general, many unexpected associations between shifts in CWMs of the complete communities and bioclimatic variables can be explained by the idiosyncratic responses of top contributors to human-induced land use changes. Top contributors drive the unexpected positive association between body mass and temperature (0.269 ± 0.0111; *p* < 0.0001) and negative associations between relative beak and wing lengths and temperature (beak: -0.243 ± 0.00827; wing: -0.387 ± 0.0106; both *p* < 0.0001; Fig. 1C). These unexpected morphological shifts can be explained by a few large-bodied species that are increasingly abundant in urban, suburban, and inhabited wildlands despite warming temperature, such as Canada Goose (*Branta canadensis*) and Wild Turkey (*Meleagris gallopavo*; Fig. 3), and species with relatively longer beaks and wings that are declining over time, such as Western Meadowlark (*Sturnella neglecta*), Horned Lark (*Eremophila alpestris*), and Barn Swallow (*Hirundo rustica*). On the other hand, relative beak and wing lengths of the top contributors are strongly positively associated with proportions of settlements (beak: 0.287 ± 0.0165; wing: 0.0836 ± 0.0187; both *p* < 0.0001), whereas these associations are strongly negative for the standard contributors (beak: - 0.278 ± 0.0174, *p* < 0.0001; wing: -0.0644 ± 0.0184, *p* = 0.000593; Fig. 1C). The positive association for relative wing sizes may arise from the increase of long-winged songbirds, such as American Robin (*Turdus migratorius*) and Red-eyed Vireo (*Vireo olivaceus*) in urban areas, while the positive association for relative beak sizes may arise from the decrease of small-beaked species such as House Sparrow (*Passer domesticus*; Fig. 3).

The unexpected associations between CWMs of demographic traits and climatic variables are again driven by the top contributors, as indicated by their responses to land use changes. One of the most important contributors to the CWM of clutch size is Northern Bobwhite (*Colinus virginianus*; Fig. 3), which has a large clutch size and has been declining precipitously due to habitat degradation (45), corresponding to stronger negative associations with agriculture and settlements of top contributors (agriculture: -0.305 ± 0.0297; settlements: -0.257 ± 0.0192; both *p* < 0.0001; Fig. 1C). The strongly positive associations between CWMs of generation length and agriculture and cultured lands are also driven by the top contributors (agriculture: 0.394 ± 0.0240; cultured land: 0.267 ± 0.0225; both *p* < 0.0001; Fig.1C), such as increasing abundances of Canada Goose and Northern Cardinal (*Cardinalis cardinalis*), both of which have above-average generation lengths (Fig. 3). Overall, our analyses provide strong evidence that changes in the abundance of a few hyper-dominant species have been responsible for driving the functional shifts in North American avian communities. Contrary to expectations that climatic conditions drive consistent morphological changes among species, these unexpected functional shifts on the community level are driven by the highly idiosyncratic responses of individual species as they benefit or perish because of human-induced changes in land cover and habitats (45–48).

At the continental scale, most of the unexpected functional shifts and their strong associations with human-induced land use changes can be explained by the top contributors, which are often stronger than, or even opposite to, the associations of CWMs calculated with the standard contributors. To test the generality of this result among local communities, we repeated the analyses on the temporal trajectories for the top contributors and standard contributors. In 1970, subcommunities composed of the top contributors were mostly characterized by having larger body mass, appendage sizes, and longer generation lengths (Fig. 2A-B). In 2021, subcommunities with top contributors have all shifted to smaller clutch sizes and longer generation lengths (Fig. 2B; fig. S3). Strikingly, spatial patterns of overall directions of functional shifts from 1970 to 2021 are nearly identical between complete communities and top contributors, indicating that the top contributors have driven most of the observed functional shifts of complete communities (Fig. 2B). In contrast, there were no widespread shifts in functional positions within the standard contributors between 1970 and 2021 (Fig. 2B; fig. S3). The uniform functional shifts of top contributors are further illustrated by the temporal decrease in spatial heterogeneity of functional positions (-0.000906 ± 0.000333; *p* = 0.00894), whereas spatial heterogeneity has increased for complete communities and standard contributors (complete: 0.00289 ± 0.000515, *p* < 0.0001; standard: 0.000605 ± 0.000267, *p* = 0.0279; Fig. 2C; table S2). Overall, our observations emphasize the hyper-dominance of a few species in driving nearly universal changes in CWMs of functional traits (49, 50), indicating that environmental drivers may not induce shared responses across all species within communities. Together, our results highlight the importance of considering the role of a few key contributors in designing conservation and management strategies. Instead of the entire community, future studies should also consider analyzing species contributions in subcommunities with specific ecological roles, such as predators, herbivores, or scavengers, to better understand the effect of global change on biotic interactions and ecosystem functioning.

## Conclusion

Rapid changes in global environments are driving within-species responses and are reshaping ecological communities, but are these community-level shifts emergent outcomes of general within-species processes? In North American avian communities, we find significant shifts in the functional traits of communities that are not driven by consistent responses among species, or by obvious climatic drivers, but instead are the outcome of the idiosyncratic rise and fall of a few highly impactful species in response to land use changes. These widespread, abundant, and often functionally distinctive species have undergone strong changes in relative abundance, leading to local and continental functional shifts that are not aligned with expectations derived from within-species trends. Our evidence highlights the challenges in predicting both within-species and community-level changes in functional traits from single environmental factors. On the other hand, our approach provides a direction in predicting community composition by focusing on the abundance and fitness of key species that contribute greatly to community structure. By analyzing summary metrics such as community-weighted means of traits essential to ecosystem functioning, we can further project the broader impact of species redistribution and identify their key contributors. Integration of functional ecology across scales will be essential to more accurately predict biotic responses to rapid global change.

## Acknowledgements

We thank Casey Youngflesh for comments on the manuscript, and members of the Weeks and Zhu labs and IGCB Postdoc Fellows for their feedback on the project. We thank Olivia Stein for creating the illustrations used in Figure 1 and 3. We thank the numerous volunteers who collected data for the North American Breeding Bird Survey. H.-X.Z was supported by the Postdoctoral Fellowship from the Institute for Global Change Biology at the University of Michigan. Y.S. was supported by the Eric and Wendy Schmidt AI in Science Postdoctoral Fellowship, a Schmidt Sciences program. K.Z. was supported by the National Science Foundation award 2306198 and USDA McIntire-Stennis Capacity Grant award 25-PAF01509. B.C.W. was supported by the David and Lucile Packard Foundation. This research was supported in part through computational resources and services provided by the Advanced Research Computing at the University of Michigan, Ann Arbor.

## Authors contributions

H.X.-Z. conceived the idea and designed the study with K.Z. and B.C.W.; H.X.-Z. curated and analyzed data with supervision from K.Z. and B.C.W. A.C.S. assisted with estimating and adjusting bird abundance; Y.S. assisted with analyzing species contributions. H.X.-Z. wrote the first draft, and all authors contributed to subsequent review and editing.

## Data and materials availability

All primary data are publicly available, and all generated data will be permanently archived upon acceptance. Code is available on GitHub: https://github.com/hengxingzou/Zouetal2025bioRXiv and will be archived upon acceptance.

## Methods

### Species abundances and trends

We estimated species relative abundance based on data from the North American Breeding Bird Survey (NABBS), a long-term survey program started in 1966 (26). We then transformed estimates of relative abundance (relative number of individuals of a given species, compared to other times and locations; not to be confused with relative proportions of species abundance in a community) into estimates of abundance (total number of individual birds, comparable across species, locations, and times) following the methods used in previous broad-scale studies of North American bird populations (38). NABBS is a structured monitoring program that has tracked changes in bird communities across much of North America, since the late 1960s. The NABBS supplies annual estimates of population change across most of Canada and the United States for almost all terrestrial bird species (with exceptions that include some waterbirds and nocturnal species; (26)). NABBS surveys are conducted each breeding season (May, June, or July) along fixed routes of 40 km at 50 evenly spaced, independent stops. At each stop, the observer counts all identifiable birds for 3 minutes within a 400 m radius. The routes are established following secondary roads, in a random design, stratified within grid-cells of one-degree latitude and longitude. NABBS routes cover a wide range of the United States and Canada, with at least one route in most grids with suitable roads.

To estimate species relative abundances, we downloaded the 2022 release that contains survey data from 1966 to 2021 from the U.S. Geological Survey website. We consider grid cells of one degree to be “local” communities; this scale represents the design stratification of the NABBS and is small enough that we do not expect significant variation of bioclimatic conditions within cells and large enough to contain enough survey data to estimate bird relative abundances (at least one NABBS route). To estimate relative abundances, we fit hierarchical Bayesian models with a spatial intrinsic conditional autoregressive structure using the R package *bbsBayes2* (51). To summarize, the model estimates relative abundance indices from NABBS counts on each route and year with a negative binomial distribution and weakly informative priors, and the model fitting of a species from shared information among all sites (“spatial strata”; in our case, one-by-one-degree grids). In addition, the model accounts for the spatial connection among strata by constructing a network graph, in which estimates in each stratum draw information from neighbors to estimate spatial variance and to assist in the model fitting (a “spatial smoothing” among neighbors). This spatial model not only allows for the estimation of relative abundances on a finer spatial scale but also outperforms the nonspatial counterpart in predicting population trajectories over time in poorly sampled locations (52). After accounting for spatial and temporal autocorrelation and variability in observation effort, the output of the model is a continuous year series of relative abundance estimates within the entire historical and current spatial range of each species. See (52) for the model structure and an expanded technical description of the model structure.

Prior to model fitting, we consolidated all records to the species level, pooling all subspecies or morphotypes and removing observations of unidentified species. The Great White Heron was combined with the Great Blue Heron (*Ardea herodias*), and the former Cordilleran, Pacific-slope, and unidentified Western Flycatchers were all pooled into Western Flycatcher (*Empidonax difficilis*). We excluded species with fewer than 50 records in the whole NABBS dataset (across all routes and years). We also excluded the White-crowned Pigeon (*Patagioenas leucocephala*), which was only recorded in one grid-cell. To constrain our analyses to high-quality data, we limited the analysis to data from 1970 to 2021, removed regions outside of the continental United States and Canada (south of 25°N, north of 55°N, and west of 135°W), and excluded local communities with fewer than 10 species in all years. Finally, we excluded nocturnal and pelagic species, as identified by the EltonTraits 1.0 dataset (16), because the NABBS protocol makes it poorly suited for estimating relative abundances for these groups. In total, we conducted analyses on 470 species in 1261 grids (Fig. 1A). We matched species taxonomy in NABBS and functional trait datasets (see below) to the 2023 version of the Clements taxonomy (15).

We used a spatially explicit, “first difference” model to estimate time series of annual relative abundance (i.e., the species’ local population trajectory) from the NABBS data for each of these 470 species. This model has been commonly used to estimate population status and trends from the NABBS data (52, 53). It estimates a species’ population trajectory in each grid as a series of annual differences from the previous year, sharing information on the estimated annual differences in a spatially explicit way (i.e., among neighboring grids using an intrinsic conditional autoregressive structure). However, we do not formally compare model performance to determine the best model structure for each species because: (1) model comparison via leave-one-out cross-validation requires strong assumptions that individual observations are conditionally independent, which is unlikely with NABBS data; (2) different models may fit better for different species, making the pooling of all species relative abundances within a local community difficult; (3) previous work comparing alternative time series models with the NABBS data has shown that the estimated population trajectories are very similar among models (51, 52).

To estimate species relative abundances, we first fit the Bayesian models of all selected species with 1000 warm-up and 1000 sampling iterations, then obtained the posterior median of relative abundance indices in each year using the *bbsBayes2* R-package. The large spatial models for each species contain thousands to tens of thousands of parameters; we therefore used a less conservative convergence criterion by considering models as converged if no parameters had Rhats > 1.1 as converged. For 31 out of 470 species for which not all the parameters converged, we reran the models, using 2000 iterations as warm up and retaining the next 2000 sampling iterations, then we closely examined the model diagnostics for the few parameters that still had not converged in 10 species’ models. The spatial model estimates parameters for each local community, enabling us to plot the maximum Rhat for each local community in the species’ range. We found that for American Robin (*Turdus migratorius*), grid cells that included a parameter with Rhat > 1.1 were from data-sparse locations outside of our study region (north of 25°N, south of 55°N, and east of 135°W), so we refit the model with only the data in our study region for 4000 sampling iterations. The rest of the species have patchy geographic distributions in some or all parts of their ranges: local communities in which species are detected are not spatially adjacent, leading to fewer neighboring communities and, therefore, poor performance of the spatial model. For these nine species, we reran the model with an alternative approach to identifying spatial neighbors (Voronoi tessellation of the grid-centers within the species’ range); this method increases the total number of neighbor relationships (i.e., broadens the definition of neighbor beyond the queen’s-rule adjacency used in the default approach; (51)), increases the sharing of information among grids, and thereby improved the model fit, especially for species with more patchy distributions (51, 52).

Finally, we transformed the relative abundances from the Bayesian models for each of the 470 species into estimates of abundance (i.e., total number of birds for each species, year, and grid-cell), based on methods used in earlier analyses (38, 54) that account for variation in each species’ detection probabilities during NABBS sampling protocols. The relative abundance of each species in each year and local community is multiplied by a series of adjustment factors that account for variations in detectability of different species. Briefly, distance adjustments account for the distances at which different species can be detected by visual or auditory signals, pair adjustments account for the probability that one individual of the breeding pair may be more easily detected (e.g., singing male vs. skulking female), and time-of-day adjustments account for the variation of detectability by the time of the survey among different species (54). NABBS was designed to compare abundance within species, not between species; these adjustments are therefore essential to our community-level analyses for not only improving accuracy but also making estimates between species comparable. We obtained the adjustment coefficients of 402 out of 470 species from (38). For the rest of the species, which are mostly waterbirds (34 species) and waterfowl (28 species), we followed previous methodologies to calculate the time-of-day adjustments (54). We then filled in missing distance and pair adjustments using the means for species within the same genus (if available), family, or order. Finally, we estimated distance and pair adjustments for Wood Stork (*Mycteria americana*), the only species without all the above information, by its behavioral patterns (no difference in detectability between sexes, and detectable from a long distance; (55)). In general, these methods remove the biases in observed counts among species by adjusting for the higher detectability of large, conspicuous, and soaring species such as pelicans, herons, and crows and the lower detectability of smaller songbirds and inconspicuous species such as warblers, wrens, and hummingbirds. We did not rerun the analyses by removing species with newly calculated adjustments because it would fundamentally change the community compositions. Rather, we checked the sensitivity by rerunning all analyses without adjustments for all species. Although these adjustments are necessary to accurately compare abundances among species, we have provided supplemental results for the full analysis without these adjustments (fig. S13-S15). Some of our conclusions are not qualitatively affected by these adjustments. In fact, our main conclusion that a small number of species greatly contribute to functional shifts across North America remained, because many common, large-bodied species that often have extreme values in functional traits are down-weighted by the adjustments. A complete species list with the adjustment factors used to estimate corrected abundances can be found in **Supplementary Data S1**.

### Functional traits, community-weighted means, and species contributions

We obtained the following morphological traits from AVONET (17): beak length (both the tip of the beak to the culmen and the tip of the beak to the nares), beak width, beak depth, the length of the relaxed wing chord, Kipp’s distance, the length from the carpal joint to the tip of the longest secondary, and body mass. We obtained data on clutch size from (56) and generation length from (36). Generation length is a composite measurement that roughly represents the average parental age (35). To obtain a holistic size metric for beak and wing while accounting for their phylogenetic correlations among species, we conducted phylogenetic principal component analyses (pPCA) separately on the four beak traits and three wing traits of the 470 species in our analyses using the R package *phytools* (57) and the dated tree from the 2023 version of Clements taxonomy (15). The first PC axes (PC1) explained 78.04% of and 86.94% of the total variance for the beak and wing traits, respectively (fig. S16). To correct for the tight correlation between beak and wing lengths, clutch size, and generation length with body mass (27, 33, 35), we conducted phylogenetic linear models for log-transformed values of each of these traits, with log-transformed body mass as the predictor variable, then extracted the residuals from each model (58). For beak and wing lengths, these residuals are considered as separate traits from beak and wing sizes, and we named them relative beak/wing lengths for the consistency with previous studies (27). Together, we calculated community-weighted means for seven functional traits: beak PC1, wing PC1, relative beak length, relative wing length, body mass, corrected clutch size, and corrected generation length.

We calculated community-weighted means (CWMs) at each year t from the relative abundance of species present in each grid (28):

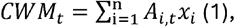

where *A*_*i,t*_ is the local relative abundance of species *i* at year *t*, calculated as the proportion of species *i*’s relative abundance at year *t* over the sum of relative abundances of all species in that community and year. *n* is the number of species in the local community (i.e.,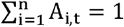), and *x*_*i*_ is the value of the trait *Q* for species *i*. We characterized the temporal trend of CWM for each trait *Q* by a simple linear model:

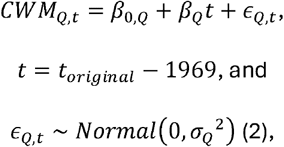

where the estimated 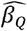 is the temporal slope, and the transformed year is the difference between the original year and 1969, such that the first year in the data (1970) was set to 1. All *t* in the following models were transformed as such.

To further partition the contributions of each species to the temporal slope of CWMs, we decomposed 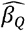 by first presenting it as the Ordinary Least Squares (OLS) solution using the variance-covariance form:

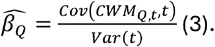

Combining Eqns. 1 and 3, 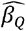 can be decomposed as

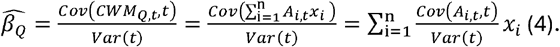

On the other hand, we can estimate the temporal slope of species *i*’s relative abundance changes as:

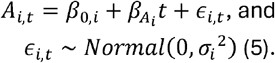

Eqn. 5 can be written with the OLS solution using the variance-covariance form as well:

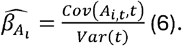

Combining Eqns. 4 and 6, the temporal slope of the CWM of trait *x* is then

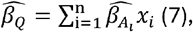

i.e., the contribution of species *i* is its temporal slope times its trait value. Note that this decomposition (Eqn. 7) is mathematically exact—the temporal trend of CWM change is exactly the summation of individual species trends (see fig. S17-S18 for an example using real data from a local community).

Using this decomposition, we calculated species contributions to temporal changes in CWMs in each local community. For each trait, we first ranked the species in each local community by absolute values of their contributions, then counted the number of local communities in which a species ranked in the top 10% in that local community. We then ranked all species by this count and pooled the top 10 species for each of the seven traits, defined as the “top contributors” (fig. S6-S12). For each local community, we conducted the following analyses using three sets of species: the full set (complete community), a subcommunity containing only the species among the top contributors, and a subcommunity containing the rest of the species (standard contributors). We also recalculated CWMs for the top contributors and the standard contributors separately for further analyses. Note that no variables (CWMs, and relative abundance) were transformed or scaled in the calculation, but time was subtracted from the starting year (see Eqn. 2). However, for better visualization, we scaled all trait values to their averages among all 470 species and used these scaled values as the x-axis of Fig. 3. We did not fit actual linear models to test for statistical significance; therefore, adjustment for multiple comparisons were not necessary. To test the sensitivity of our analyses, we conducted all analyses with the top five instead of top 10 contributor for each trait and found no qualitative changes in our main results (fig. S19-20).

### Temporal shifts in the functional space

To further characterize the temporal change of local communities, we mapped the community in a grid each year in a functional space shared among all communities over years, constructed with a PCA of CWMs of the five key traits (beak PC1, wing PC1, body mass, generation length, and clutch size). We then tracked the location of each local community over time within the functional space, creating a trajectory, and characterized the community trajectory using the position of their start and end points (1970 and 2021) and the overall direction of change from 1970 to 2021. The overall direction (effective angle) was measured as the angle relative to the positive x-axis of the vector pointing from the beginning (1970) to the end (2021) of the trajectory (Fig. 3A). Finally, we constructed and analyzed trajectories from the CWMs of the two subcommunities: top contributors and standard contributors. To ensure all trajectories were comparable, we projected the subcommunities onto the PC axes of the PCA with the CWMs of the complete community so that all trajectories share the same axes. After obtaining the position of each local community in the functional space, measured in quadrants, we spatially mapped the local community each year and calculated the spatial heterogeneity by Moran’s *I* statistic using the *spdep* package (59). We compared the time series of spatial heterogeneity calculated from the CWMs of the base community and the two subcommunities to further illustrate the contribution of the top 23 species to the spatial patterns of functional shift over time.

To compare patterns of functional shifts across major biogeographical regions, we categorized each local community into its bird conservation regions (BCRs; (60)). For spatial grids that fall into multiple BCRs, we calculated the percentage area in each BCR and only considered the one with the highest percentage. We further generalized the BCRs in the continental U.S. and Canada into seven major “avifaunal biomes” (61): Arctic, Northern Forest, Pacific, Intermountain West, Southwest, Prairie, and Eastern (fig. S5).

### Environmental data

We quantified the association between shifts in CWMs and changes in key environmental variables to explore the potential mechanisms of community-level functional shifts and to facilitate their comparisons with within-species expectations. We obtained a time series of the mean temperature and precipitation in the warmest quarter of the year (*bio*10 and *bio*18) from the WorldClim2 dataset (62). The warmest quarter coincides with both the breeding season of most North American birds and the period when NABBS is conducted. Both *bio*10 and *bio*18 are strongly correlated with mean annual temperature (MAT; linear model, slope = 0.650, 95% CI = [0.649, 0.651], *p* < 0.0001) and mean annual precipitation (MAP; slope = 0.173, 95% CI = [0.172, 0.174], *p* < 0.0001); therefore, we do not expect results to be qualitatively different if using MAT and MAP instead. To evaluate the effect of seasonality on key demographic traits, we calculated the seasonality of temperature and precipitation as the difference between the warmest and coldest months (*bio*5 *- bio*6) and the wettest and driest months (*bio*13 *- bio*14; (63)). To explore the relationship between human-induced land use change and functional shifts, we obtained land cover data from the HYDE 3.3 database, which estimates historical anthropogenic land use changes (64). The HYDE data contain 20 categories of anthropogenic biomes, or “anthromes” (9, 30); we generalized land use types into four main categories: settlements, which contains urban areas and villages; agriculture, which contains croplands and rangelands; cultured land, which contains woodland and drylands with human inhabitants, and wild land. Settlements and agriculture are intensely impacted by human activities, cultured land receives less human impact, and wildland contains woodland and drylands with minimum human impacts. To match the spatial resolution of estimated species abundances, we calculated the proportions of areas in each land use category in a one-degree latitude and longitude grid cell. We obtained yearly time series of all the above environmental data from 1970 to 2021 and aggregated them to the same one-degree grid cells as our abundance data.

### Model fitting

Before fitting the full model, we first fit the following linear mixed-effect models with the R package *lme4* (65):

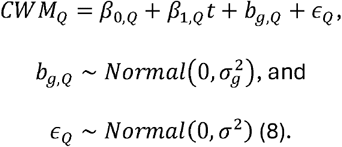

Specifically, we separately modeled the CWM of each trait *Q* as a linear response of transformed year *t* with slope *β*_1,*Q*_, intercept *β*_0,*Q*_, grid-level random effect *b*_*g,Q*_, and the overall residual *ϵ*_*Q*_. We used a simple grid-level random effect because there was no spatial autocorrelation in the residuals of the full model (see below). Both *b*_*g,Q*_ and *ϵ*_*Q*_ are normally distributed. We weighted the fitted residual 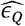 by the inverse of the overall abundance of the local community within each grid. This weighting corrects for the potentially higher measurement errors in communities with lower overall abundance. In all linear mixed effects models, we log-transformed the CWM of body mass. Because we fit the CWM of each trait separately, we adjusted the p-value of the models using the Benjamini-Hochberg method (66). Fitting the year-only model yielded significant slopes for CWMs of all traits but generation length (table S3).

Then, to explore whether the year effect still accounted for variances unexplained by all environmental variables (bioclimatic variables and human-induced land use proportions), we fit a year-environment model:

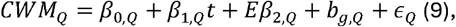

where *E* is a vector of environmental variables, *β*_2,*Q*_ is a vector of slopes corresponding to each environmental variable, and *b*_*g,Q*_ and *ϵ*_*Q*_ follow definitions in Eqn. 8. Because the land use type is proportional data (summing to 1), we removed wild land when fitting the model. We then scaled all the environmental variables to ensure the slopes are comparable in magnitude. In this model, year slopes for CWMs of all traits were significant and had the same signs comparing to the year-only model (**Supplementary Data S3**), indicating that year explained variations in CWMs of most traits even after accounting for all environmental variables, which changed temporally, and should be considered as a fixed effect rather than a temporal random effect.

Finally, we fit the full model with year, environmental variables, year-environment interactions, and grid-level random effects:

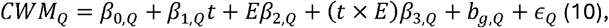

where *t* × *E* denotes the interactive effects between year and environmental variables. We analyzed the models of each CWM using variograms in the R package *gstat* (67) and found no evidence of strong spatial autocorrelation in the residuals (fig. S21). We then fit the full model (Eqn. 10) with CWMs of the two subcommunities, consisting of the top contributors and the standard contributors, then compared their slopes of the same predictor variables. We fit separate models rather than using group-level slopes within a single model because CWMs do not contain species identities and therefore can only be calculated for different groups of species. See **Supplementary Data S4** for full diagnostics of the models.

Temperature and precipitation seasonality may only affect the life history strategies of residents who are exposed to climatic extremes throughout the whole year. Therefore, we analyzed the associations between temperature/precipitation seasonality and the two demographic traits, clutch size and generation length, separately for residents and migrants. To do so, we filtered species by their migration status according to (38), then separately calculated the CWMs for each group in local communities.

### Software

All analyses were conducted by R version 4.4.1. (68). In addition to specific packages mentioned in methods, all data wrangling and visualization were conducted using the package *tidyverse* (69), and all geospatial analyses were conducted using the package *sf* (70).

