## Supplementary Material for "Hyper-dominant species drive unexpected functional shifts in North American birds"

**List of Supplementary Materials**

Tables S1-S3

Figs. S1 to S21

Supplementary Data S1-S4

**Supplemental Tables**

**Table S1.** Associations between CWMs of all traits and environmental factors, measured by linear mixed effects models. All predictor variables (environmental variables and, for clutch size and generation length, log-transformed body mass) were scaled, and CWMs of body mass, clutch size, and generation length were log-transformed. Each local community (“complete” community) is divided by the species that are the top contributors of continental-wide patterns (top contributors, or “top”) and the rest of the species (standard contributors, or “standard”). Year was transformed by subtracting 1969 such that the first year in the time series (1970) is 1 when fitting the model. For simplicity, standard errors are omitted, and all significant slopes (*p* < 0.05) are in bold. See full statistics, including estimates of interactive effects, in **Supplementary Data S2**.

| **Variable** | **Beak PC1** | **Wing PC1** | **Relative beak length** | **Relative wing length** | **Body Mass** | **Clutch Size** | **Generation Length** | **Community** |
| --- | --- | --- | --- | --- | --- | --- | --- | --- |
| Year | **5.83E-04** | **0.00198** | **0.00721** | **0.00947** | **0.0183** | **-0.00561** | **-5.79E-04** | Complete |
| Temperature of warmest quarter | **0.0353** | **0.0295** | **-0.177** | **-0.326** | **0.168** | **0.290** | 0.00597 |  |
| Precipitation of warmest quarter | **0.0136** | **-0.00390** | **-0.0191** | **-0.172** | **-0.0271** | **0.0155** | 0.00427 |  |
| Temperature seasonality | **0.00351** | -7.65E-04 | **-0.0168** | **0.0515** | **-0.0960** | **-0.112** | **0.0339** |  |
| Precipitation seasonality | **-0.00638** | **-0.00818** | **0.0563** | **-0.0504** | **-0.0315** | **0.0756** | **-0.0243** |  |
| Settlements | 0.00318 | **0.0148** | **0.0326** | -0.00416 | **0.111** | **-0.128** | **0.395** |  |
| Agriculture | **-0.0688** | **-0.0414** | **-0.153** | 0.00489 | 0.0388 | **-0.272** | **0.373** |  |
| Cultured land | **-0.0280** | -0.00168 | 0.00234 | **0.103** | **0.180** | **-0.244** | **0.331** |  |
| Year | **0.00177** | **0.00438** | **0.00291** | **0.0110** | **0.0274** | **-0.00843** | **-0.00239** | Top |
| Temperature of warmest quarter | **0.0522** | **0.0423** | **-0.243** | **-0.387** | **0.269** | **0.367** | **-0.0758** |  |
| Precipitation of warmest quarter | **0.0163** | **-0.00576** | **-0.0139** | **-0.114** | **-0.0702** | **0.183** | **0.0366** |  |
| Temperature seasonality | **-0.0142** | **-0.0117** | **0.0293** | **0.0688** | **-0.168** | **-0.132** | **0.0583** |  |
| Precipitation seasonality | -0.00162 | **-0.00993** | 0.00368 | **-0.0632** | **-0.0387** | **0.0280** | **-0.0359** |  |
| Settlements | **-0.0175** | **-0.0201** | **0.287** | **0.0836** | 0.0207 | **-0.257** | **0.379** |  |
| Agriculture | -0.0115 | **-0.0237** | 0.0468 | **0.123** | 0.0584 | **-0.305** | **0.394** |  |
| Cultured land | 0.00362 | 0.00216 | 0.0358 | 0.0346 | **0.154** | -0.0396 | **0.267** |  |
| Year | **-5.96E-05** | **5.58E-04** | **0.00714** | **0.00296** | **0.00778** | **-0.00170** | **0.00622** | Rest |
| Temperature of warmest quarter | **0.0170** | **0.0126** | **0.0256** | **-0.104** | **0.0368** | **0.140** | 0.0135 |  |
| Precipitation of warmest quarter | **0.0112** | 0.00191 | **-0.0166** | **-0.109** | **0.0182** | **-0.124** | **-0.0219** |  |
| Temperature seasonality | **0.0149** | **0.0122** | **-0.0786** | **0.0291** | **-0.0118** | **-0.0750** | **0.0221** |  |
| Precipitation seasonality | **-0.00753** | **-0.00494** | **0.0515** | **-0.0171** | **-0.0164** | **0.0655** | -0.00347 |  |
| Settlements | **0.0221** | **0.0357** | **-0.278** | **-0.0644** | **0.187** | **0.0696** | -0.0189 |  |
| Agriculture | **-0.106** | **-0.0578** | **-0.157** | **-0.0731** | **-0.0934** | 0.0206 | **-0.0979** |  |
| Cultured land | **-0.0491** | **-0.0214** | -0.0114 | **0.0669** | **0.0563** | **-0.123** | **0.151** |  |

**Table S2.** Temporal trends of spatial heterogeneity, calculated by Moran’s *I* (see Methods), for the base and the two subcommunities. Trends were estimated using linear regression.

| **Community** | **Slope** | **Standard error** | **t-value** | ***p*** |
| --- | --- | --- | --- | --- |
| Complete | 0.00289 | 5.15E-04 | 5.61 | <0.001 |
| Top | -9.06E-04 | 3.33E-04 | -2.72 | 0.00894 |
| Standard | 6.05E-04 | 2.67E-04 | 2.26 | 0.0279 |

**Table S3.** Temporal trends of all community-weighted means (CWMs) of each grid, estimated by linear regression, measured by linear mixed effects models. All predictor variables (environmental variables and, for clutch size and generation length, log-transformed body mass) were scaled, and CWMs of body mass, clutch size, and generation length were log-transformed. Each local community (“complete” community) is divided by the species that are the top contributors of continental-wide patterns (top contributors, or “top”) and the rest of the species (standard contributors, or “standard”). Year was transformed by subtracting 1969 such that the first year in the time series (1970) is 1 when fitting the model.

| **Trait** | **Slope** | **Standard error** | **Df** | **t-value** | ***p*** | **Community** |
| --- | --- | --- | --- | --- | --- | --- |
| Beak PC1 | 6.30E-04 | 1.72E-05 | 64312.2945 | 36.7304541 | <0.0001 | Complete |
| Wing PC1 | 0.00202742 | 2.00E-05 | 64312.7427 | 101.434268 | <0.0001 |  |
| Relative beak length | 0.00741304 | 8.56E-05 | 64312.074 | 86.6458654 | <0.0001 |  |
| Relative wing length | 0.00921329 | 1.30E-04 | 64316.1455 | 70.9417628 | <0.0001 |  |
| Body mass | 0.01910854 | 8.97E-05 | 64313.6401 | 213.107297 | <0.0001 |  |
| Clutch size | -0.0058921 | 1.07E-04 | 64309.0318 | -55.157003 | <0.0001 |  |
| Generation length | 1.33E-04 | 7.82E-05 | 64312.3585 | 1.70205274 | 0.0956 |  |
| Beak PC1 | 0.00134389 | 2.51E-05 | 64222.7267 | 53.602603 | <0.0001 | Top |
| Wing PC1 | 0.00416479 | 3.04E-05 | 64215.368 | 136.828916 | <0.0001 |  |
| Relative beak length | 0.00331979 | 8.25E-05 | 64310.1852 | 40.2303891 | <0.0001 |  |
| Relative wing length | 0.01332814 | 1.18E-04 | 64306.5252 | 112.560546 | <0.0001 |  |
| Body mass | 0.02706026 | 1.22E-04 | 64302.3664 | 221.806061 | <0.0001 |  |
| Clutch size | -0.0113158 | 1.24E-04 | 64287.8472 | -91.001595 | <0.0001 |  |
| Generation length | -0.0024194 | 7.71E-05 | 64306.5308 | -31.375397 | <0.0001 |  |
| Beak PC1 | 1.39E-04 | 1.83E-05 | 64312.6346 | 7.59164416 | <0.0001 | Standard |
| Wing PC1 | 6.48E-04 | 2.02E-05 | 64313.1312 | 32.05161 | <0.0001 |  |
| Relative beak length | 0.00717459 | 8.79E-05 | 64313.293 | 81.5887515 | <0.0001 |  |
| Relative wing length | 0.00191538 | 9.61E-05 | 64313.2488 | 19.9353386 | <0.0001 |  |
| Body mass | 0.00860069 | 7.41E-05 | 64313.3787 | 116.059401 | <0.0001 |  |
| Clutch size | -6.66E-04 | 9.38E-05 | 64312.0994 | -7.1055292 | <0.0001 |  |
| Generation length | 0.00525916 | 7.37E-05 | 64311.4107 | 71.3381914 | <0.0001 |  |

**Supplemental Figures**

**Figure S1.** Associations between community-weighted means (CWMs) of seven key functional traits and main environmental variables, including climate, human-induced land use, and year, calculated from resident species only. Blue and orange dots represent negative and positive associations, respectively.

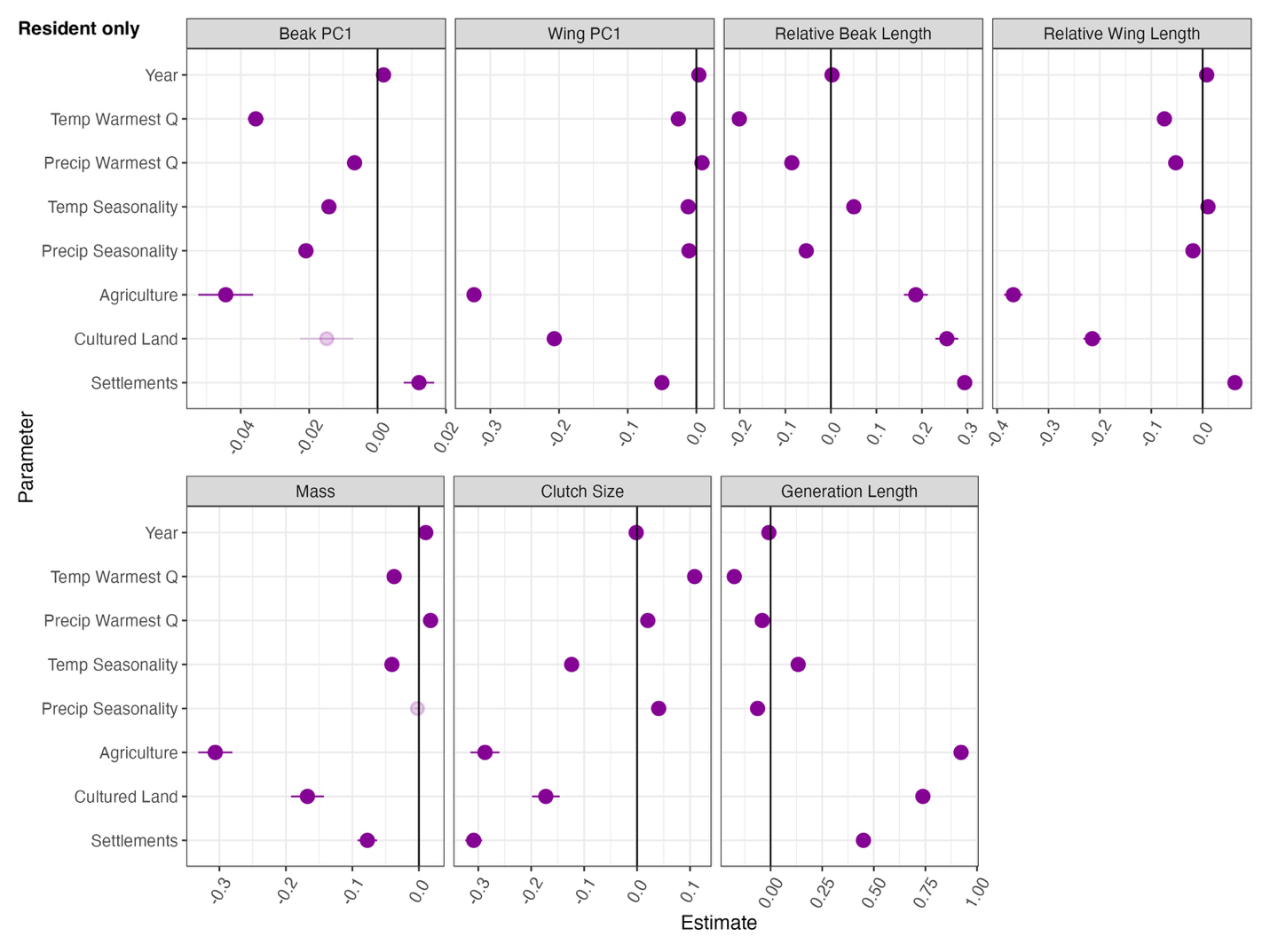

**Figure S2.** Associations between community-weighted means (CWMs) of seven key functional traits and main environmental variables, including climate, human-induced land use, and year, calculated from migratory species only. Blue and orange dots represent negative and positive associations, respectively.

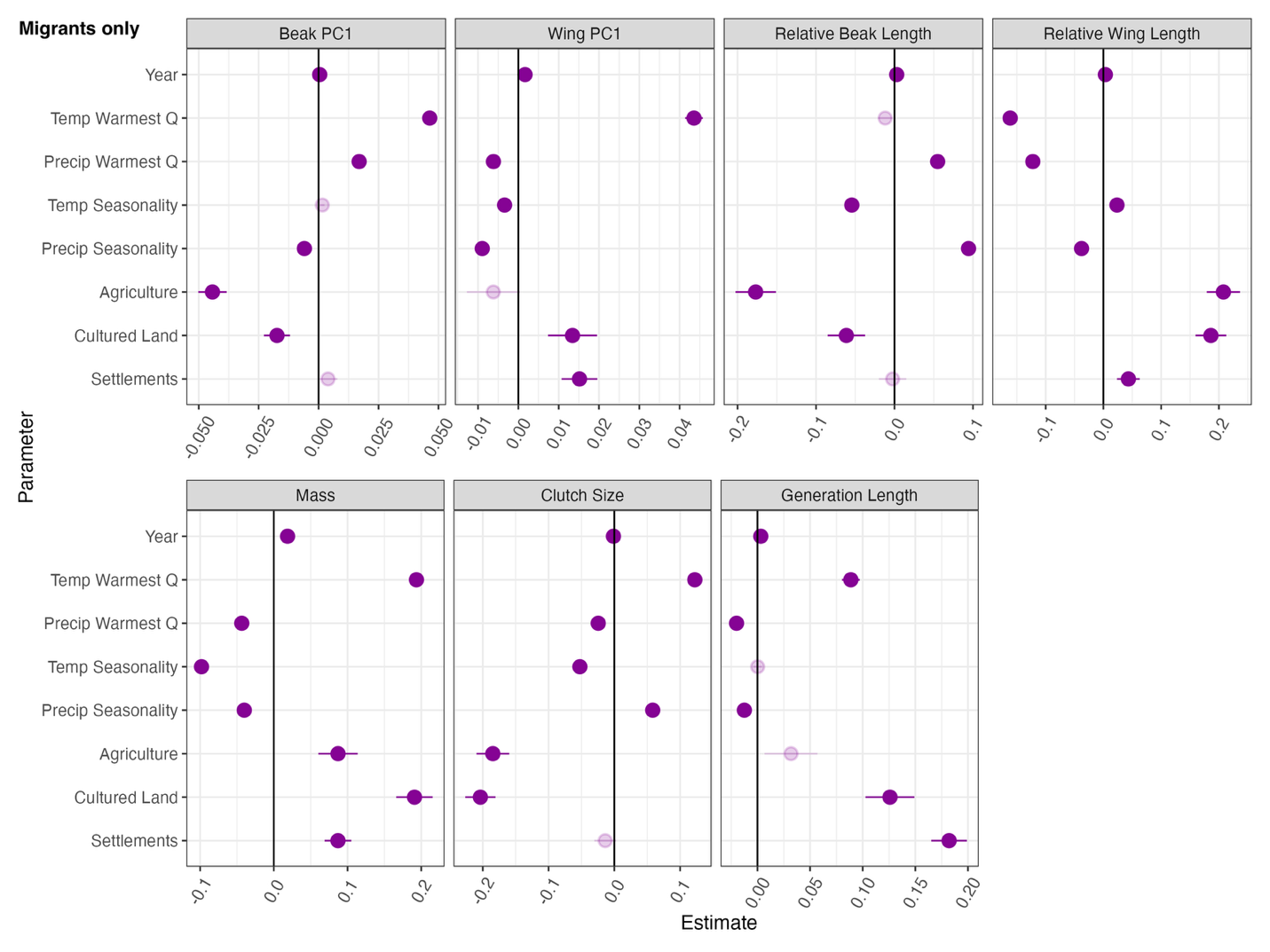

**Figure S3.** Proportions of functional positions (start and end points) and functional shifts (effective angles) across North America; see Fig. 2A for definitions of the functional positions and shifts.
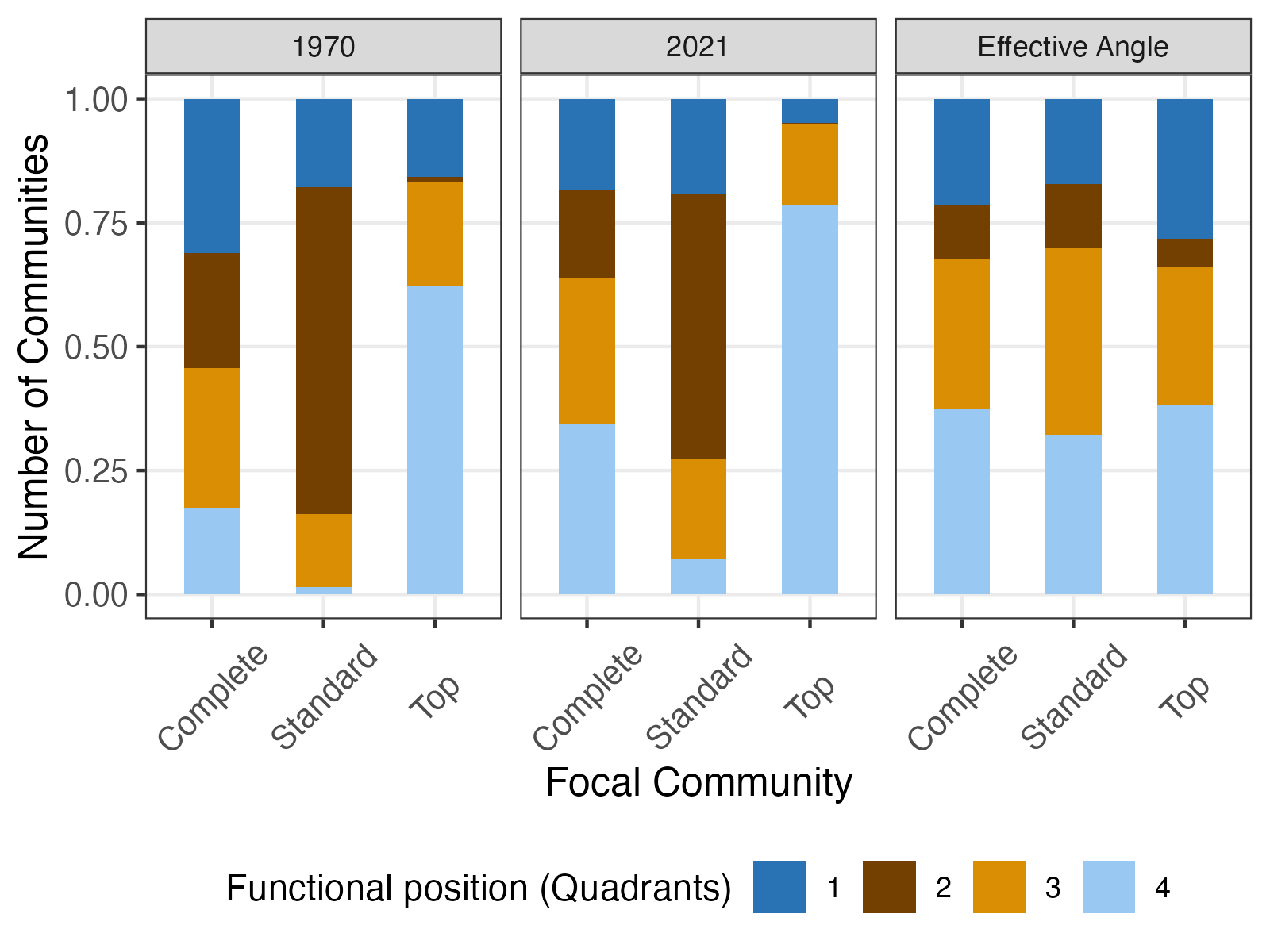

**Figure S4.** Proportions of functional positions (start and end points) and functional shifts (effective angles) across North America by avian biomes. See Fig. 2A for definitions of the functional positions and shifts; see fig. S5 for a map of major avian biomes.
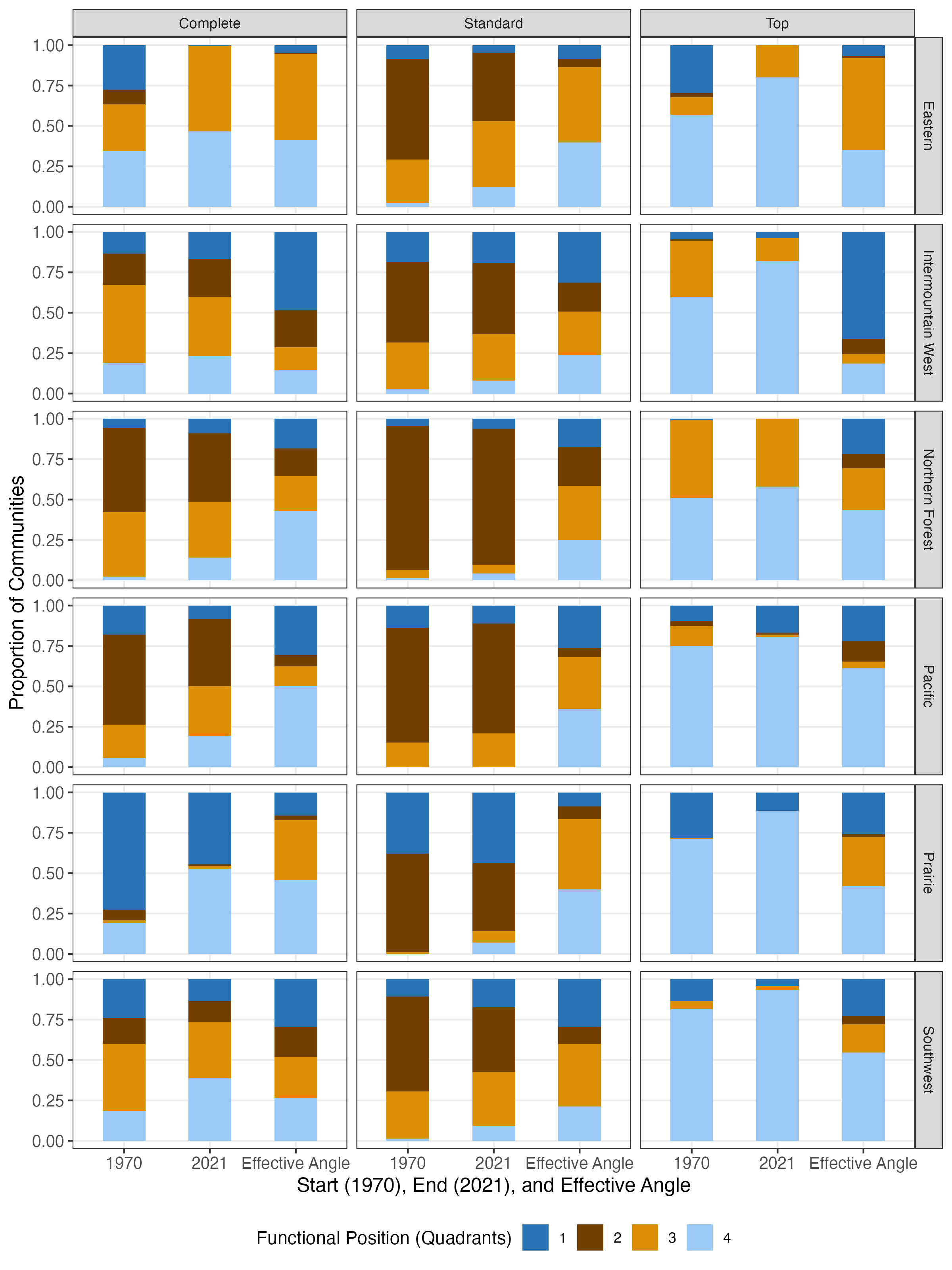

**Figure S5.** Map of avian biomes in Contiguous United States and Canada.
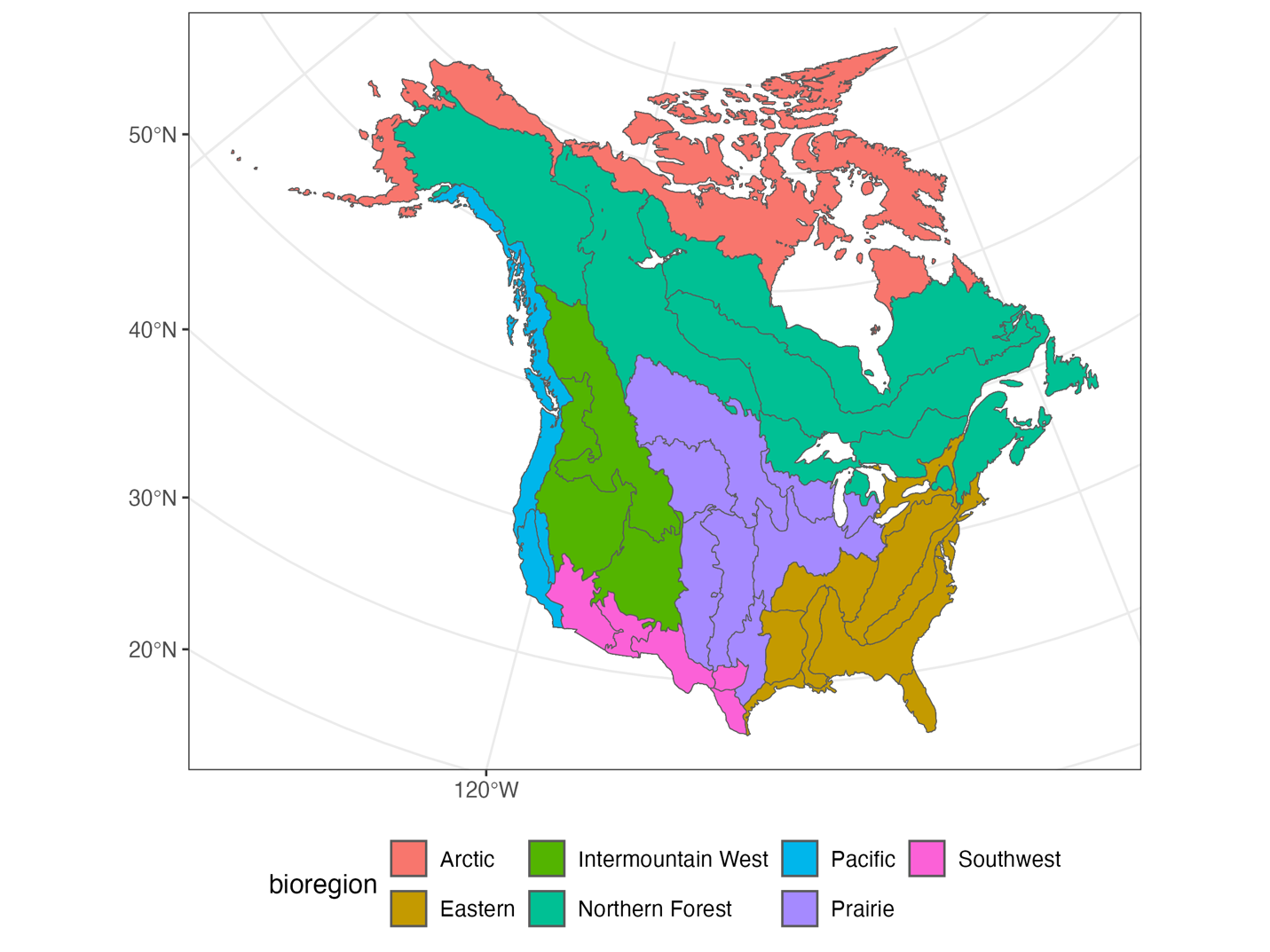

**Figure S6.** Top 10 contributors to the temporal trends in the CWM of beak size.

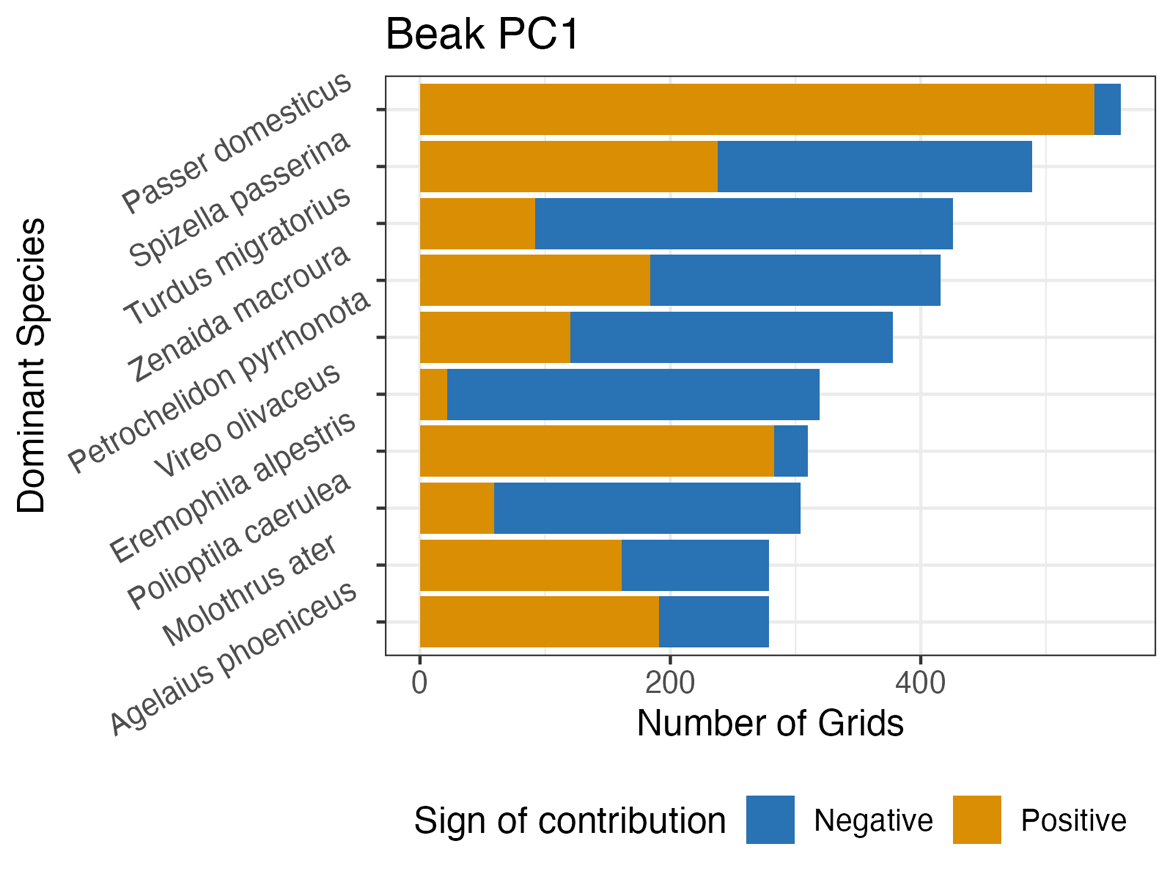

**Figure S7.** Top 10 contributors to the temporal trends in the CWM of wing size.

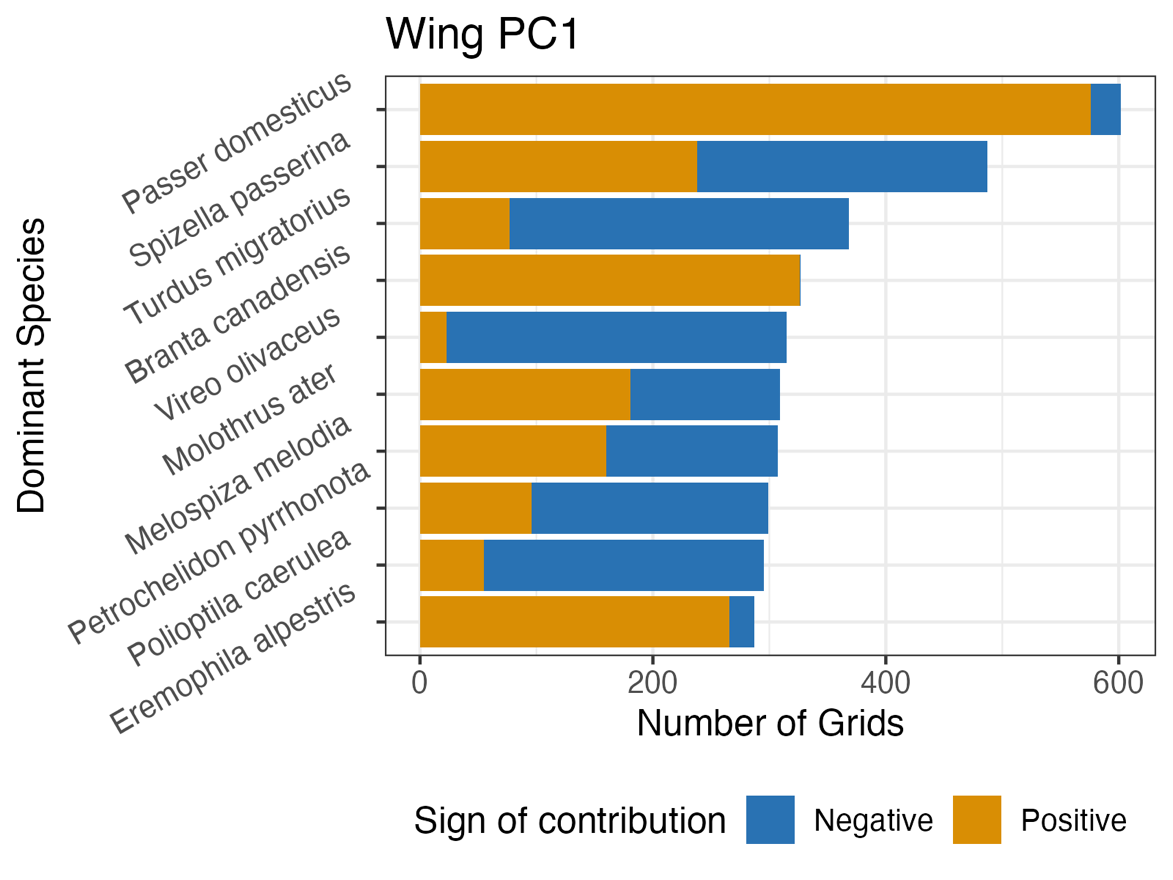

**Figure S8.** Top 10 contributors to the temporal trends in the CWM of relative beak length.

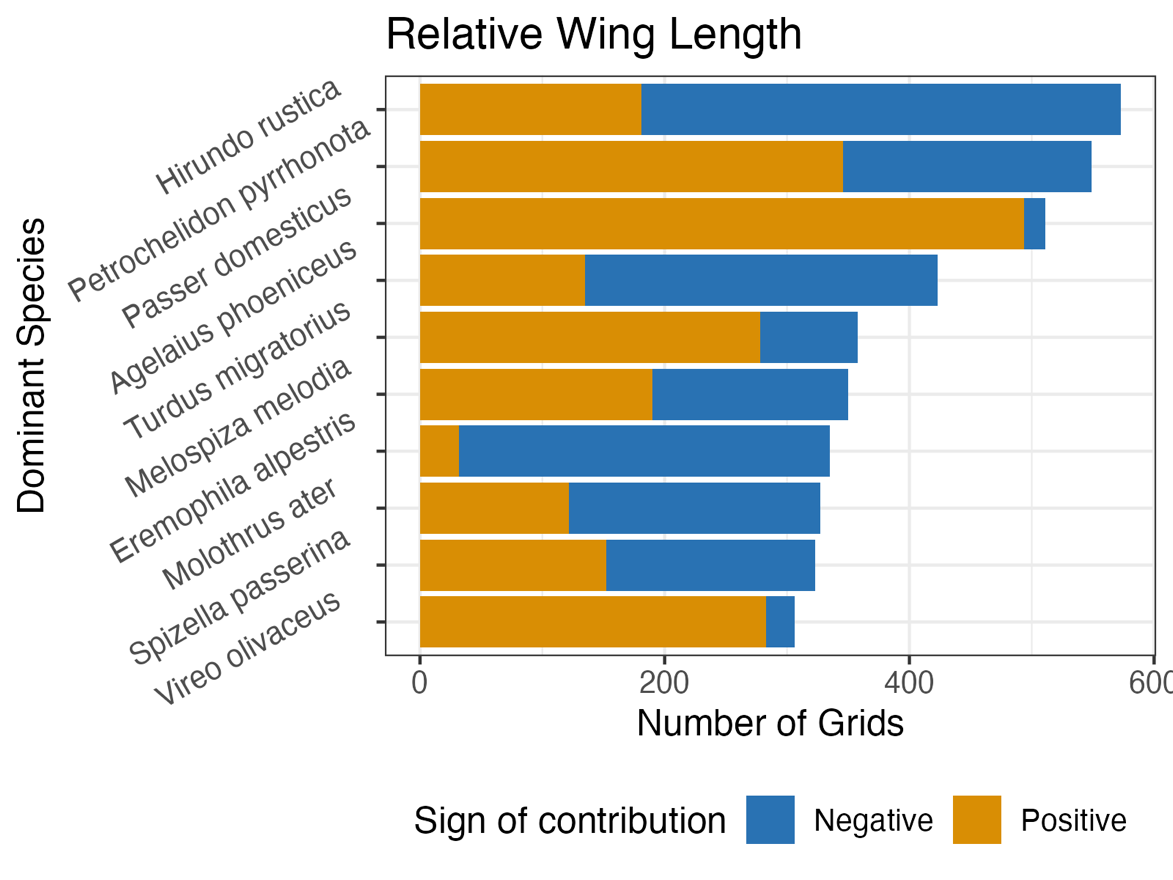

**Figure S9.** Top 10 contributors to the temporal trends in the CWM of relative wing length.

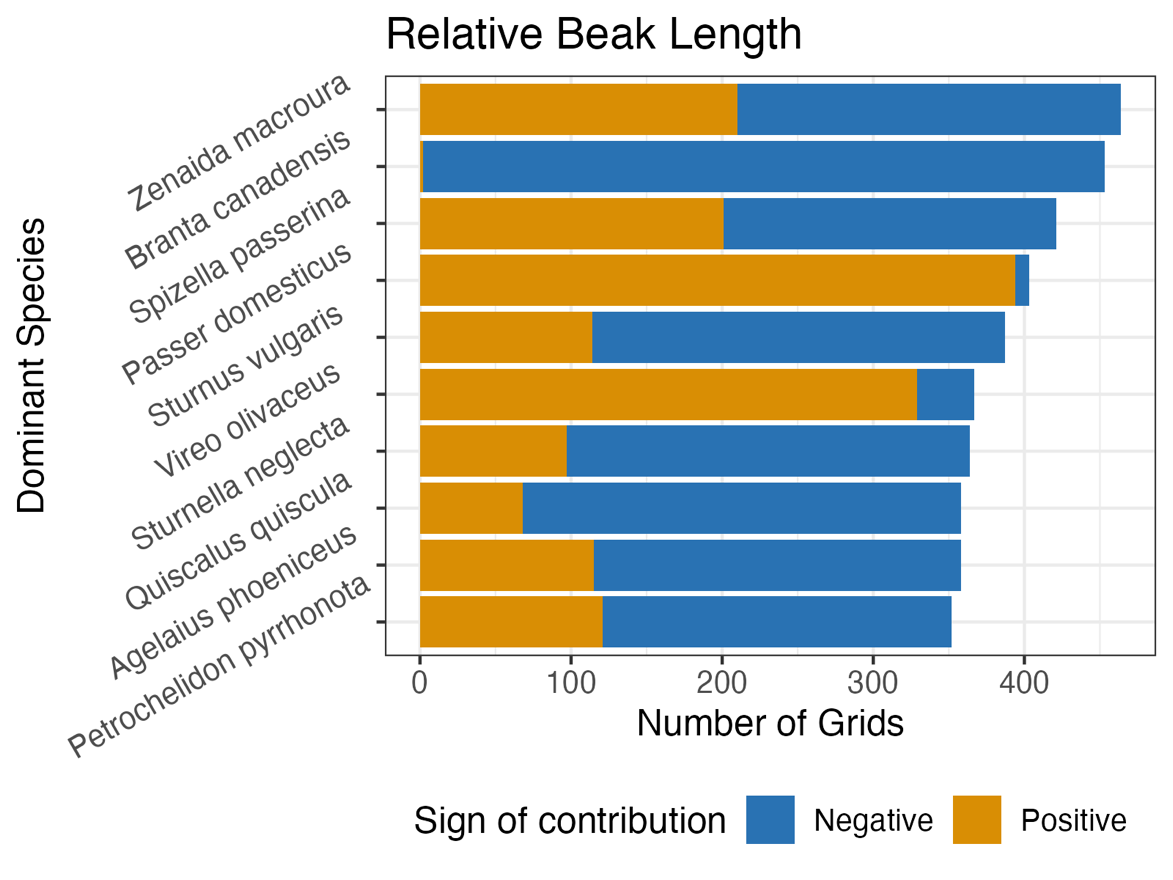

**Figure S10.** Top 10 contributors to the temporal trends in the CWM of body mass.

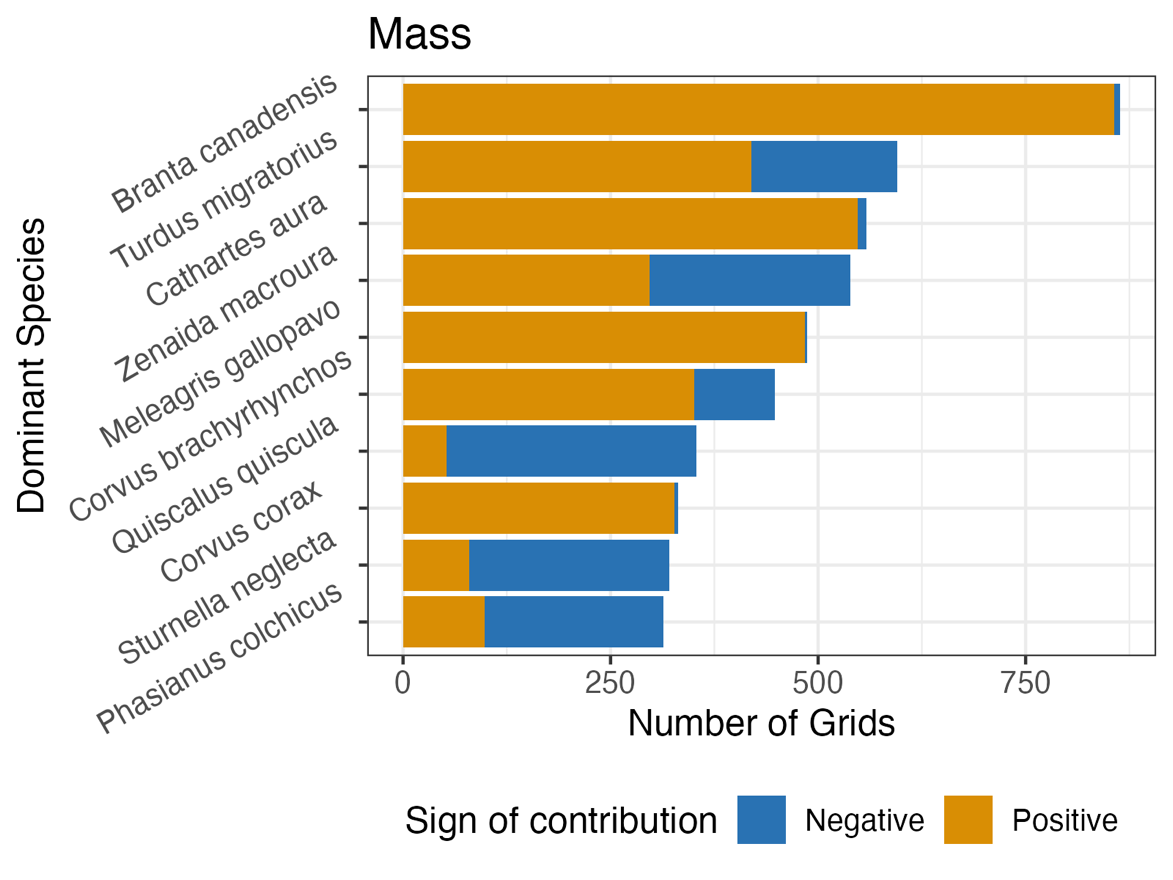

**Figure S11.** Top 10 contributors to the temporal trends in the CWM of clutch size.

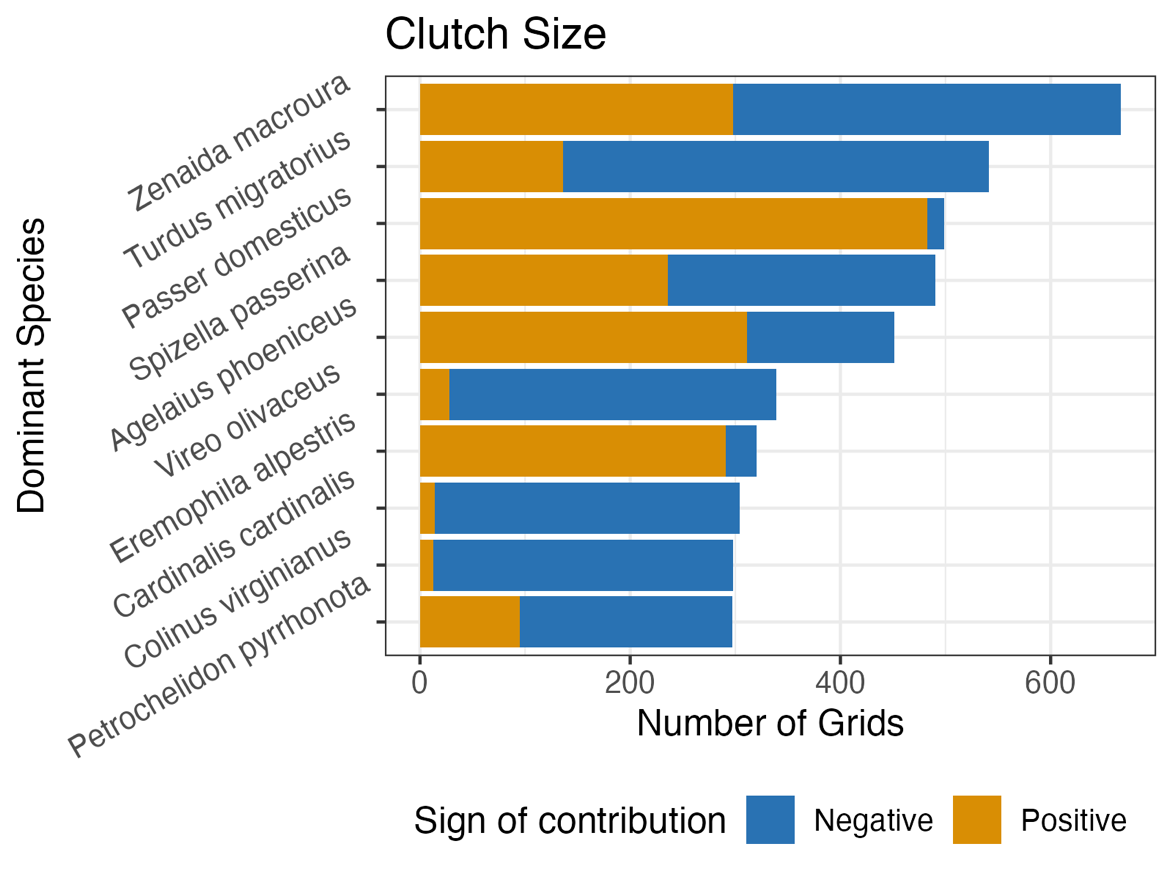

**Figure S12.** Top 10 contributors to the temporal trends in the CWM of corrected generation length.

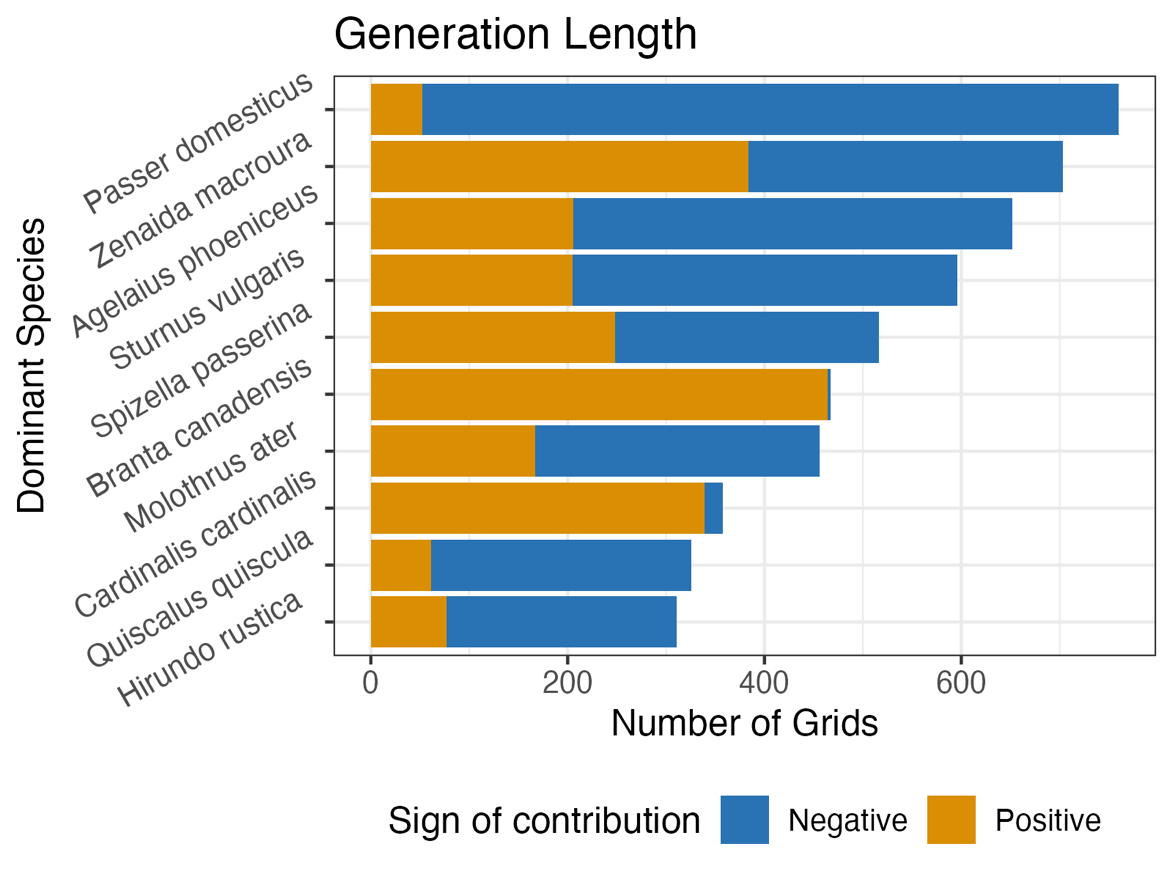

**Figure S13.** Results generated without correcting species abundances with detectability. Associations between community-weighted means (CWMs) of seven key functional traits and main environmental variables, including climate, human-induced land use, and year. CWMs are calculated from either all species in the communities (complete communities), only the top contributors, or only the rest of the species (standard contributors). Compare to Fig. 1A.

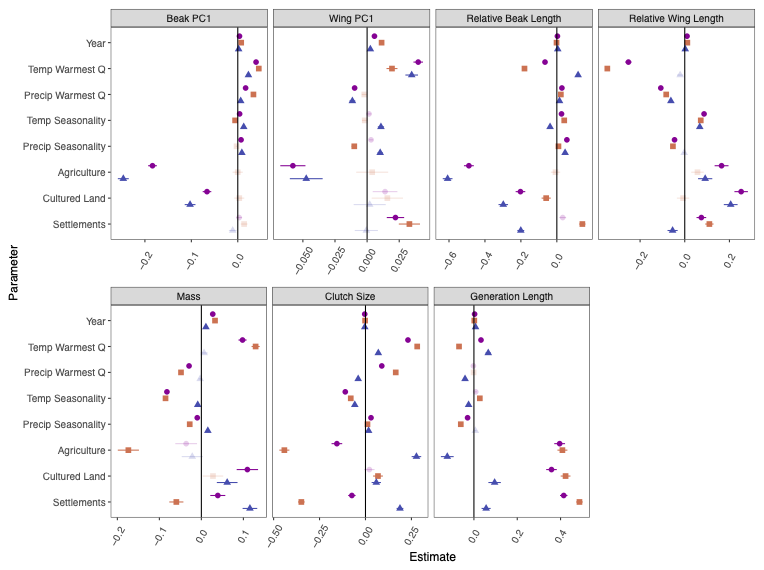

**Figure S14.** Results generated without correcting species abundances with detectability. **A.** The functional space constructed by PCA of all communities by their community-weighted mean measurements over time, and a temporal trajectory of one local community (40˚N, 110˚W) moving through the functional space over time, showing the start and end points and the overall direction (effective angle). In the trajectory, lighter shades represent earlier years, and darker shades represent later years. The two panels represent the same functional space, but the scales are adjusted for better visualization. **B.** Maps of each local community’s start and end positions in the functional space, measured by the quadrants in **A**, and the overall direction of the functional shifts, measured by the quadrants to which the trajectory travels. Columns show results for complete communities, subcommunities with top contributors, and subcommunities with the standard contributors. **C.** Spatial heterogeneity of functional positions of the base communities, top contributors, and the rest of the species, calculated as Moran’s *I* from the maps. A Moran’s *I* closer to 0 means lower spatial heterogeneity. Compare to Fig. 2.

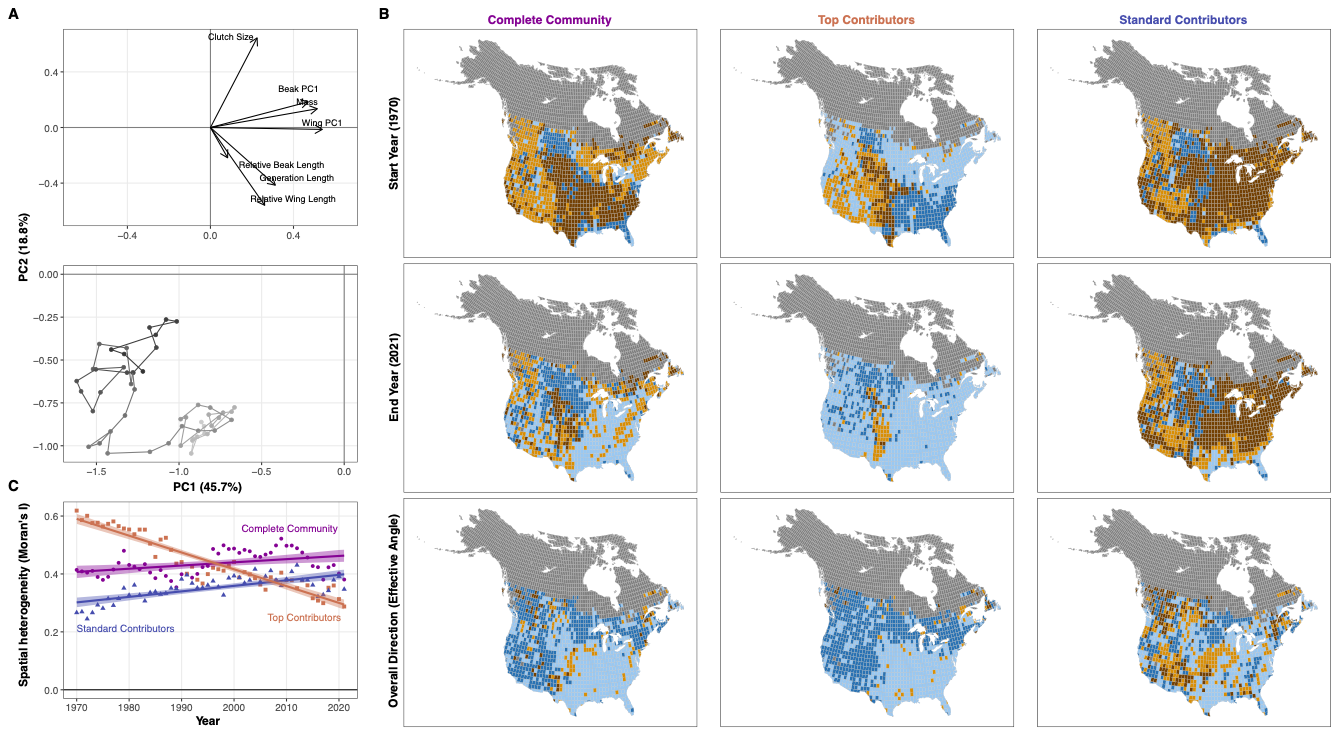

**Figure S15.** Species contributions to the community-weighted means (CWMs) of seven key functional traits. Panels only include species that contribute substantially (ranked top 10% of all species in a local community) in at least one local community; point size denotes the number of grids in which a species contributes greatly. The sign of the contribution depends on the species’ temporal trends of relative abundance and its trait value, e.g., if a species with below-average trait value increases in relative abundance, it will contribute negatively by lowering the CWM of the trait. For each trait, the identity of the top 10 species that contribute to the CWMs of the most local communities is labeled; pooling all these species results in 25 top contributors of the observed functional shifts over time, shown on the right. For each panel, the x-axis is the scaled trait values, and the y-axis is the median of the temporal slopes of relative abundances across each species’ range. Shades denote the absolute magnitudes of species contributions: colder colors represent negative contributions, and warmer colors represent positive contributions. Axes and scales are different for each panel for better visualization. Compare to Fig. 3.

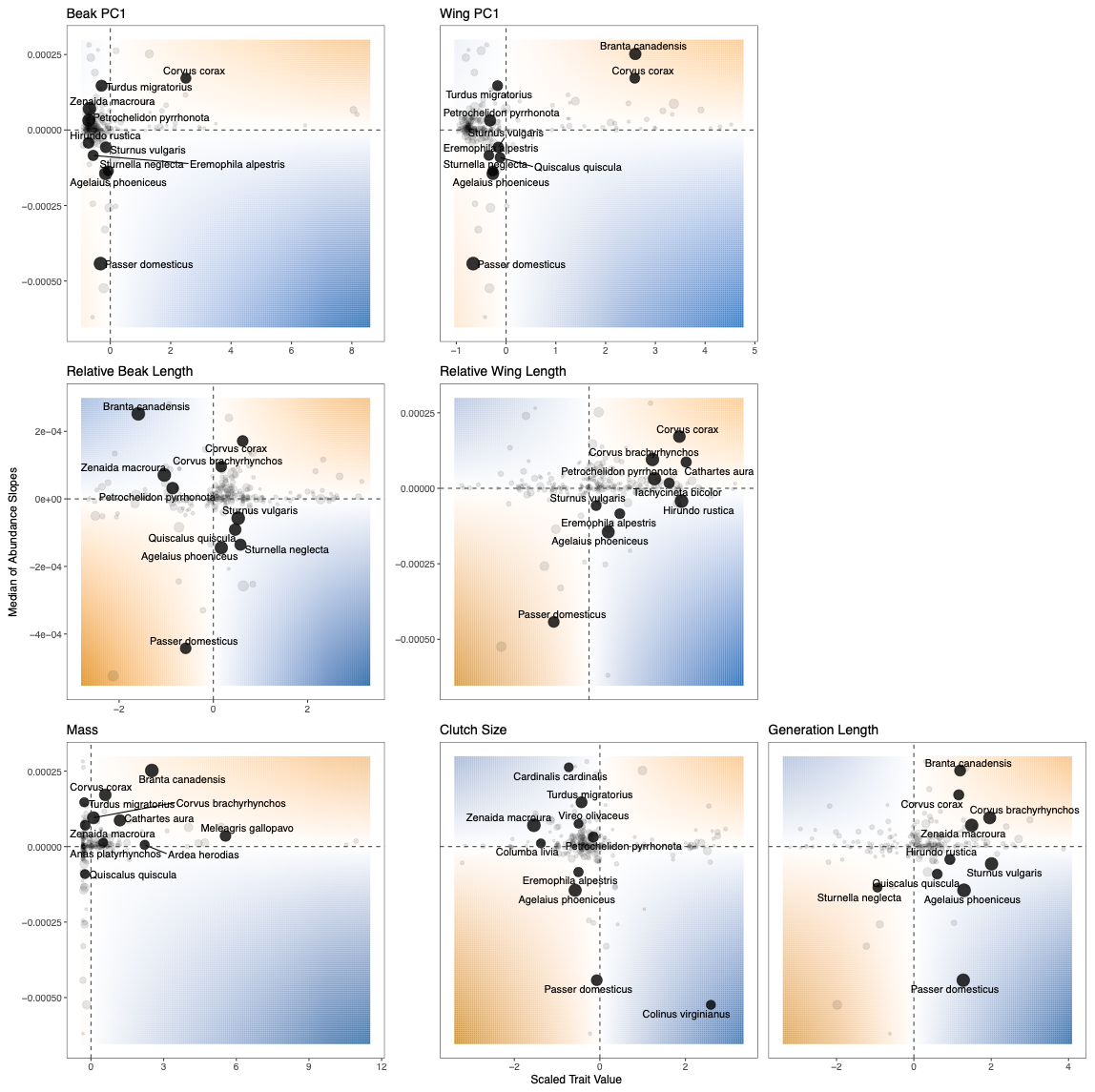

**Figure S16.** Phylogenetic principal component analysis (pPCA) of four beak (left) and three wing (right) traits. The first principal axes (PC1) of each set of traits are used to characterize beak and wing sizes.
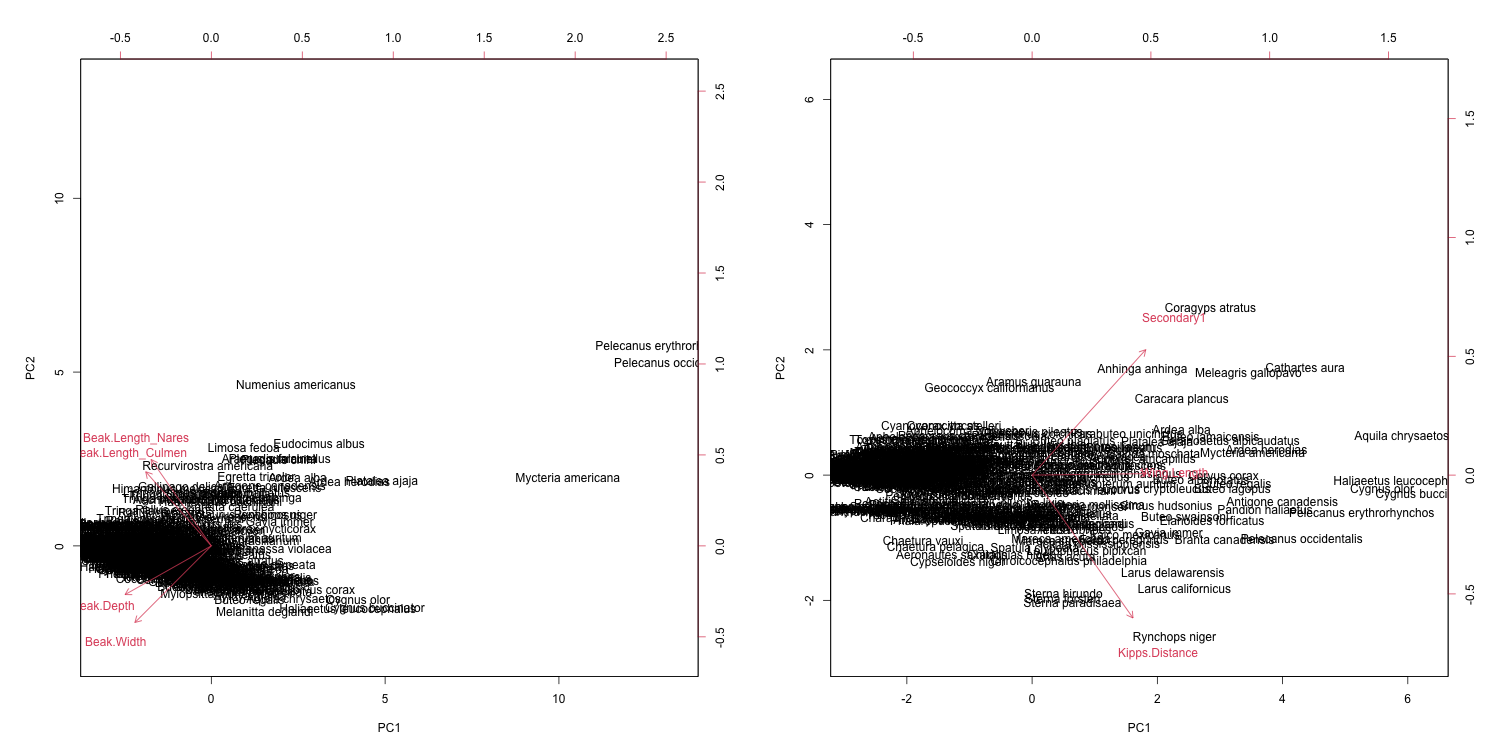

**Figure S17.** Distributions of species contributions to temporal trends of CWMs in an example local community (40˚N, 110˚W). The x-axis represents the absolute values of species contributions. For all CWMs, species contributions show a consistently skewed distribution, suggesting that only a few species contribute greatly to temporal trends of CWMs.
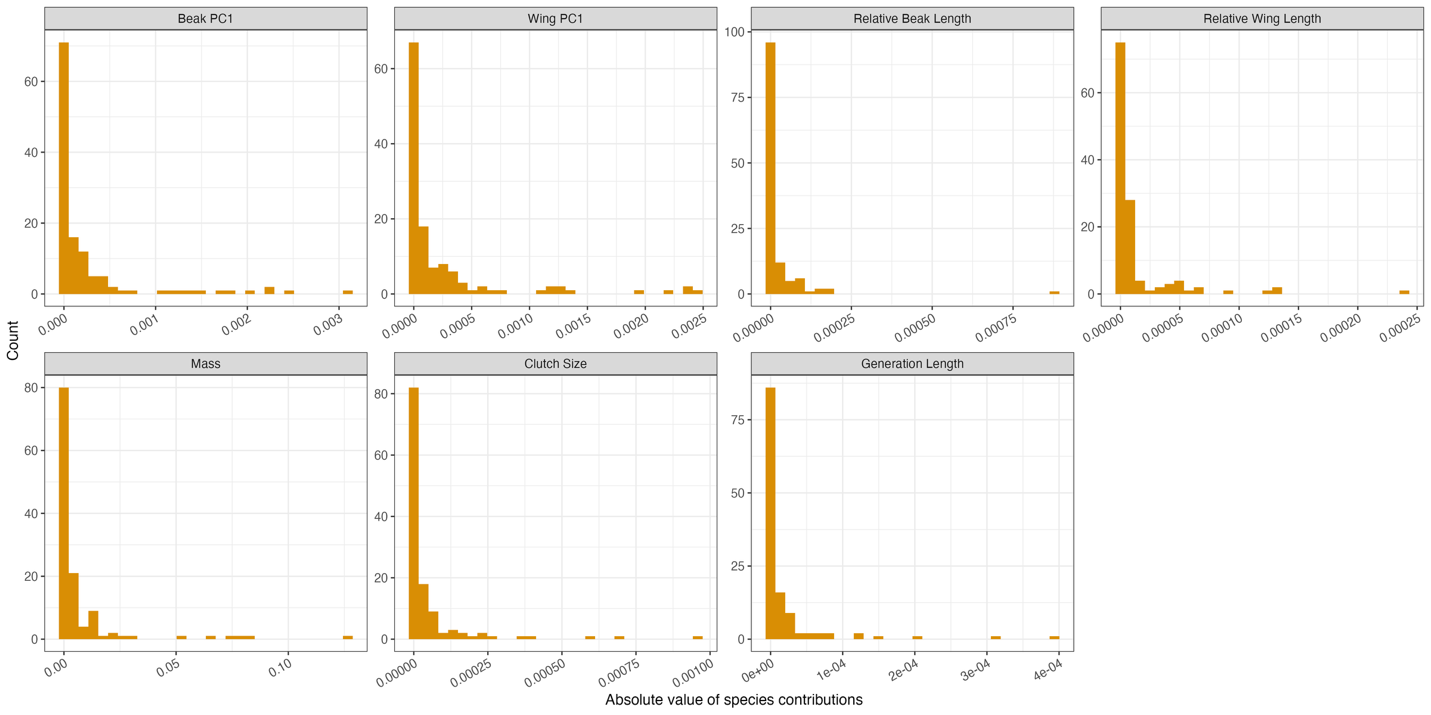

**Figure S18.** A visual example of how contributions of each species add up to the temporal slopes of CWMs, from an example local community (40˚N, 110˚W). The x-axis shows species, as ordered by their absolute values of contributions (larger on the left). The y-axis shows the cumulative contributions of all species to the left of each species. As more species are counted (i.e., towards the right of the x-axis), the cumulative contributions approach the overall temporal trends of CWMs, as shown by the horizontal orange line.
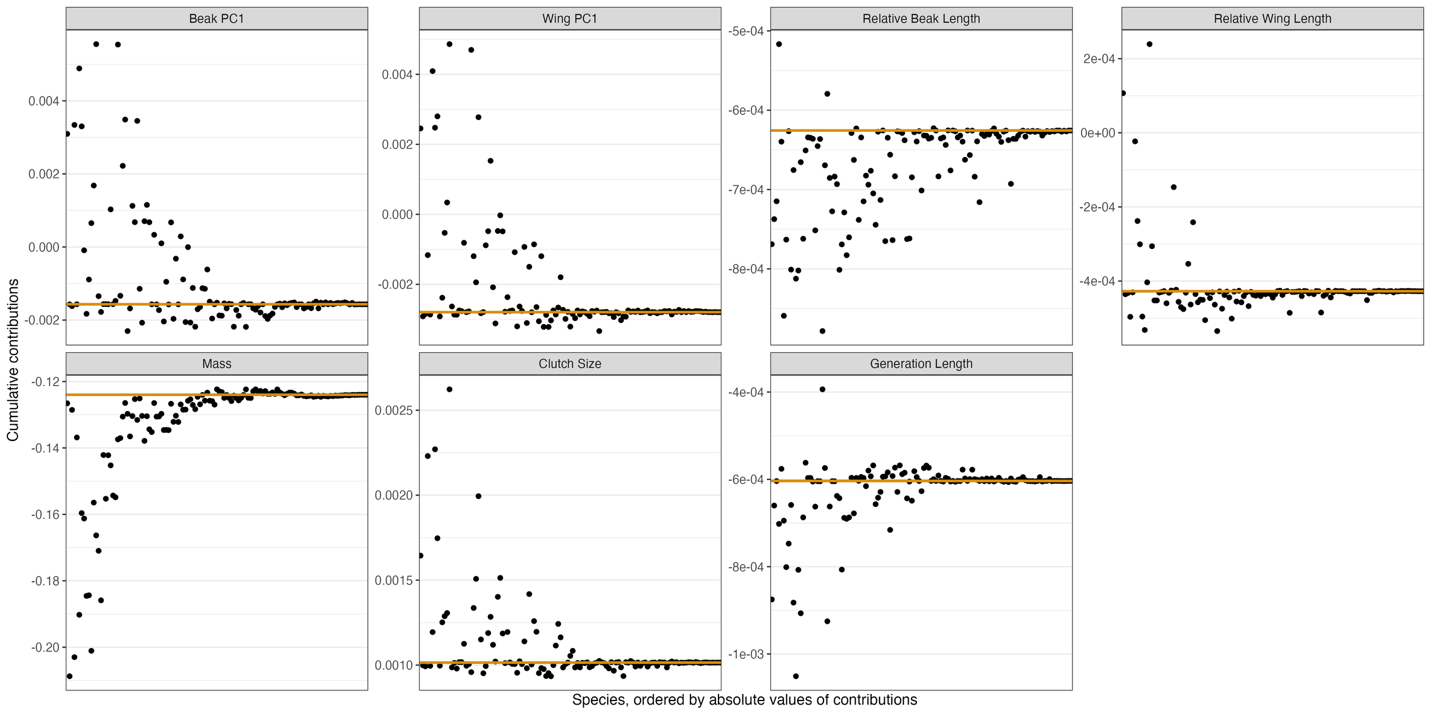

**Figure S19.** Results generated with top five instead of 10 species that contribute greatly to the CWMs of each trait. Associations between community-weighted means (CWMs) of seven key functional traits and main environmental variables, including climate, human-induced land use, and year. CWMs are calculated from either all species in the communities (complete communities), only the top contributors, or only the rest of the species (standard contributors). Compare to Fig. 1A.

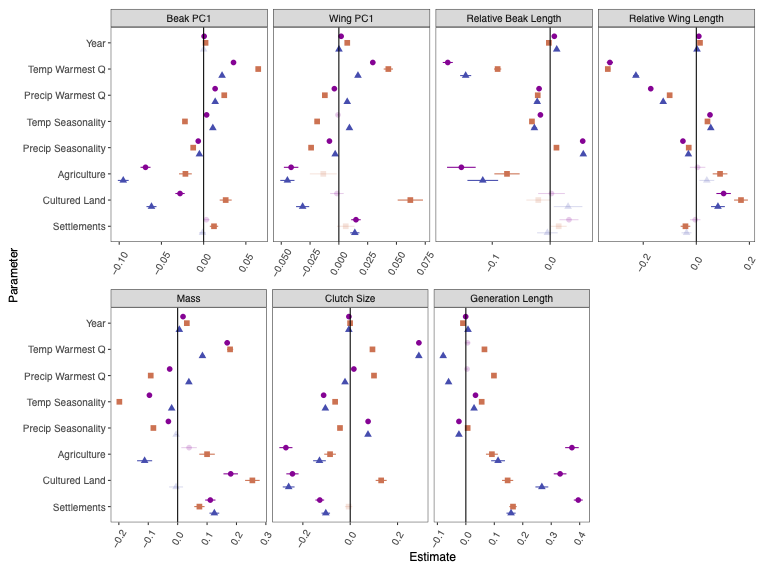

**Figure S20.** Results generated with top five instead of 10 species that contribute greatly to the CWMs of each trait. **A.** The functional space constructed by PCA of all communities by their community-weighted mean measurements over time, and a temporal trajectory of one local community (40˚N, 110˚W) moving through the functional space over time, showing the start and end points and the overall direction (effective angle). In the trajectory, lighter shades represent earlier years, and darker shades represent later years. The two panels represent the same functional space, but the scales are adjusted for better visualization. **B.** Maps of each local community’s start and end positions in the functional space, measured by the quadrants in **A**, and the overall direction of the functional shifts, measured by the quadrants to which the trajectory travels. Columns show results for complete communities, subcommunities with top contributors, and subcommunities with the standard contributors. **C.** Spatial heterogeneity of functional positions of the base communities, top contributors, and the rest of the species, calculated as Moran’s *I* from the maps. A Moran’s *I* closer to 0 means lower spatial heterogeneity. Compare to Fig. 2.

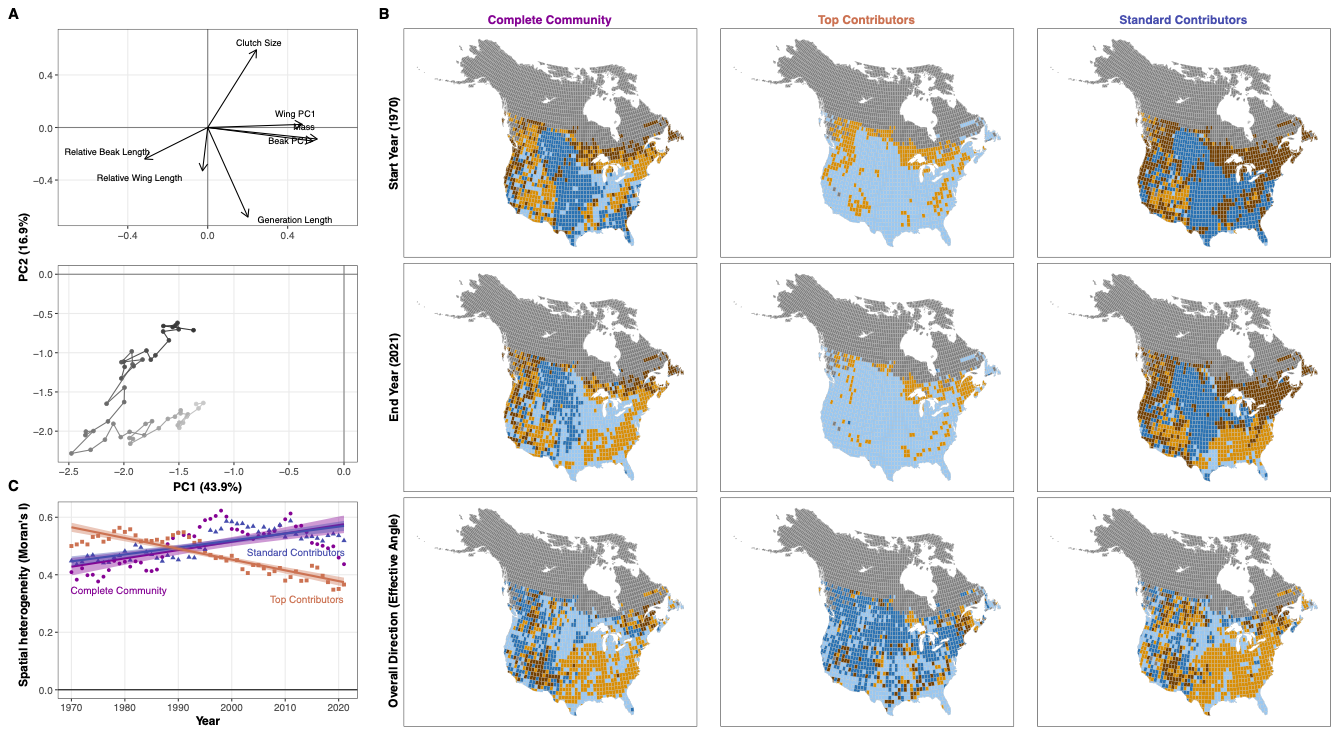

**Figure S21.** Variogram of the linear mixed effects models for each of the seven functional traits, with CWMs calculated from the complete communities. The x-axis indicates the distance, and the y-axis indicates the semivariance in the residuals of the model. A strong spatial signal in the residual (i.e., spatial autocorrelation unaccounted for in the model) would lead to an increase in the semivariance, then plateauing at a certain value. In our models, none of the traits clearly show this pattern, indicating that there is no or little spatial autocorrelation in the residuals.

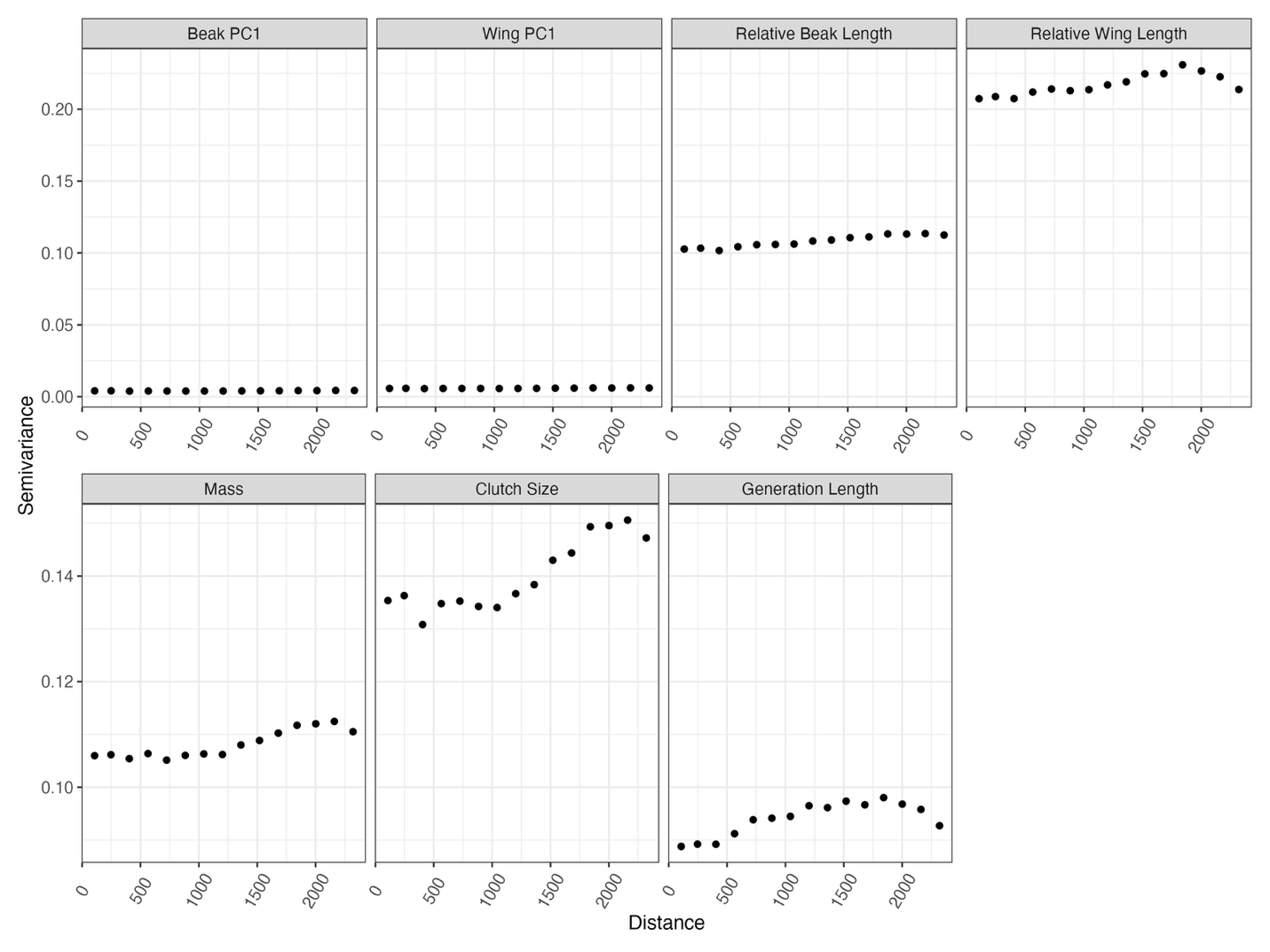
